# Recent secondary contact, genome-wide admixture, and limited introgression of neo-sex chromosomes between two Pacific island bird species

**DOI:** 10.1101/2023.09.13.557611

**Authors:** Elsie H. Shogren, Jason M. Sardell, Christina A. Muirhead, Emiliano Martí, Robert G. Moyle, Daven C. Presgraves, J. Albert C. Uy

## Abstract

Secondary contact between closely related taxa represents a “moment of truth” for speciation. Removal of geographic barriers allows us to test the strength of reproductive isolation that evolved in allopatry and identify the genetic, behavioral, and/or ecological barriers that separate species in sympatry. Sex chromosomes are known to rapidly accumulate differences between species, an effect that may be exacerbated for neo-sex chromosomes because they are regions of the genome that have recently become linked to sex chromosomes and are transitioning from autosomal to sex-specific inheritance. Two closely related bird species in the honeyeater family — *Myzomela cardinalis* and *Myzomela tristrami* — carry neo-sex chromosomes and have come into recent secondary contact in the Solomon Islands after being isolated for ∼3 my. Hybrids of these two species have been observed in sympatry for at least 100 years. To determine the genetic consequences of hybridization, we use population genomic analyses of individuals sampled in allopatry and sympatry to characterize gene flow in the contact zone. Using genome-wide estimates of diversity, differentiation, and divergence, we find that the degree and direction of introgression varies dramatically across the genome. Autosomal introgression is bidirectional, with phenotypic hybrids and phenotypic parentals of both species showing admixed ancestry. On the sex and neo-sex chromosomes, the story is different. Introgression of Z is limited and neo-Z sequence shows no evidence of introgression, whereas introgression of W and neo-W is strong but highly asymmetric, moving only from the invasive *M. cardinalis* to the resident *M. tristrami*. Thus, reproductive isolation is incomplete, but sex and neo-sex chromosome regions have prevented gene flow in one (W/neo-W) or both (Z/neo-Z) directions. The recent contact between previously isolated species indicates that hybridization may permit gene flow between taxa in some genomic regions, but species divergence can be maintained by barriers to gene flow associated with rapidly evolving sex-linked regions of the genome.

**Author Summary:** When a new species colonizes an island and interacts with a closely related native species, we are provided with a rare opportunity to identify the factors that keep species distinct and the consequences of interbreeding. Regions of the genome that evolve rapidly or influence mate choice may be especially likely to act as barriers to gene flow. The red *Myzomela cardinalis,* birds in the honeyeater family, have recently arrived to Makira in the Solomon Islands, joining the endemic, all black *Myzomela tristrami.* We used population genomic analyses of individuals in geographic isolation, as well as those in geographic contact with the other species, to understand the history of these two species and the consequences of their recent range overlap on Makira. We found that regions of the genome that are sex-specific (*i.e.,* sex chromosomes) were either limited in their ability to move between species, or only moved in one direction, from the invading *M. cardinalis* to the native *M. tristrami*. This work highlights how certain regions of the genome may be especially important in defining species boundaries and the generation and maintenance of biodiversity.

## Introduction

When taxa are geographically separated, it is difficult to test whether two groups are good biological species that no longer freely interbreed with each other [1–3]. Secondary contact, however, provides an opportunity to test the strength of reproductive isolation in sympatry and was considered the “moment of truth” for speciation [1,4]. The presence or absence of interspecific pairings and/or hybrid offspring serve as phenotypic proxies for reproductive isolation, but with genetic data we now know that hybridization between seemingly good biological species is not uncommon (reviewed by [5]). A more genic view of speciation allows us to consider which regions of the genome are most important in defining and maintaining species divergence [6]. Typically, secondary contact is studied in long-standing hybrid or tension zones [7,8], in which the interaction of gene flow, selection, and recombination occurring over hundreds to thousands of generations allows specific loci to be identified that either move between species or act as reproductive barriers [9–13]. However, characterizing patterns of introgression in the earliest stages of range overlap may be useful in identifying the regions of the genome that are primary barriers to gene flow, influencing patterns of introgression before evolution in sympatry can occur. Identifying the extent and direction of admixture in recently sympatric taxa can therefore be informative about the genetic and/or phenotypic factors which either facilitate or prevent gene flow [14,15].

Introgression can occur if traits under sexual or natural selection are globally adaptive, increasing fitness in the genomic background of the either species [16,17]. Neutral alleles may also introgress due to the demographic dynamics of range expansion precipitating the contact event [18]. However, gene flow can be limited by locally adaptive alleles which result in sexual, ecological, and/or intrinsic genetic incompatibilities. Sexual incompatibilities may result from differences in courtship signals or mate preferences [1,19,20]. Ecological incompatibilities can result if hybrids possess intermediate phenotypes poorly suited to either parental habitat [21,22]. Finally, intrinsic genetic incompatibilities can reduce the fertility or viability of hybrid individuals [23–25].

Incompatibilities may be especially likely to arise on sex chromosomes, as these regions of the genome are expected to diverge rapidly. Sex chromosomes have lower effective population sizes compared to autosomes and in the heterogametic sex, recessive mutations are immediately exposed to selection, leading to a “faster-X” (or −Z) effect [26–29]. Although demography, mating system, and dosage compensation may mediate the strength of faster-X [30], empirical evidence confirms that sex-linked regions show elevated substitution rates and rapid divergence of gene expression [28,31,32]. The rapid evolution occurring on sex chromosomes can lead to them playing a disproportionately large role in speciation [33,34]. Species recognition [35] and mating behaviors [36] have also been mapped to sex-linked loci, suggesting these regions can be important in maintaining species boundaries via sexual incompatibilities [37]. Sex chromosomes are also known to limit gene flow between taxa through genetic incompatibilities that reduce hybrid viability of the heterogametic sex (Haldane’s rule; [33,38]) and/or an excess of hybrid sterility factors (large X/Z effect; [33,39,40]).

Neo-sex chromosomes — often formed by the fusion of a former autosome to an ancestral sex chromosome — are regions of the genome that are newly sex-specific, becoming heterogametic in one sex and homogametic in the other [41–44]. Accelerated evolution during the transition from diploidy could enrich neo-sex chromosomes for sexual, ecological, and/or genetic incompatibilities, which reduce gene flow between taxa [45–48]. In birds, sex chromosomes were thought to be syntenic [49,50]. However, evidence of neo-sex chromosome evolution in birds has increased dramatically in recent years; fusion of autosomal regions to sex chromosomes has been discovered in several different lineages [41,44,51–57]. Despite the potential for incompatibilities, to our knowledge only two studies have considered the role of neo-sex chromosomes in reproductive isolation in avian taxa, looking at intra-specific introgression between subspecies [58] and mitolineages [59]. Although neo-sex chromosomes are important in speciation for taxa ranging from plants to fish [43,45,48], much remains to be understood about how these genomic regions influence gene flow between species with incomplete reproductive isolation.

*Myzomela* honeyeaters of the Solomon Islands present an opportunity to study the evolution of reproductive isolation in a system with neo-sex chromosomes [4,60,61]. The sexually monomorphic, all black Sooty honeyeater (*M. tristrami*) is endemic to the large island of Makira (Fig 1A). The sexually dimorphic, red Cardinal honeyeater (*M. cardinalis*) shared a common ancestor with *M. tristrami ∼*3 mya [62]; it is found across many islands of the South Pacific, including the small satellite islands of Ugi and Three Sisters, which are 8 km and 20 km from Makira, respectively (Fig 1A). Secondary contact occurred when *M. cardinalis* expanded its range and established a narrow, coastal region of sympatry with *M. tristrami*. Intermediate plumaged birds collected on Makira in 1908 were identified as putative hybrids [63] and in 1927, Mayr collected phenotypically pure *M. cardinalis* as well as putative hybrids as part of the Whitney South Seas Expedition [64]. Subsequent expeditions did not collect any phenotypic hybrids and *M. cardinalis* was more abundant than the native *M. tristrami* in sympatry [63], leading Mayr and Diamond [4] to propose that hybridization occurred only when conspecific mates were scarce. The first genetic investigation of the system uncovered evidence for hybridization using six nuclear and two mitochondrial markers, and for potential neo-sex chromosomes [60,61].

**Figure 1.**
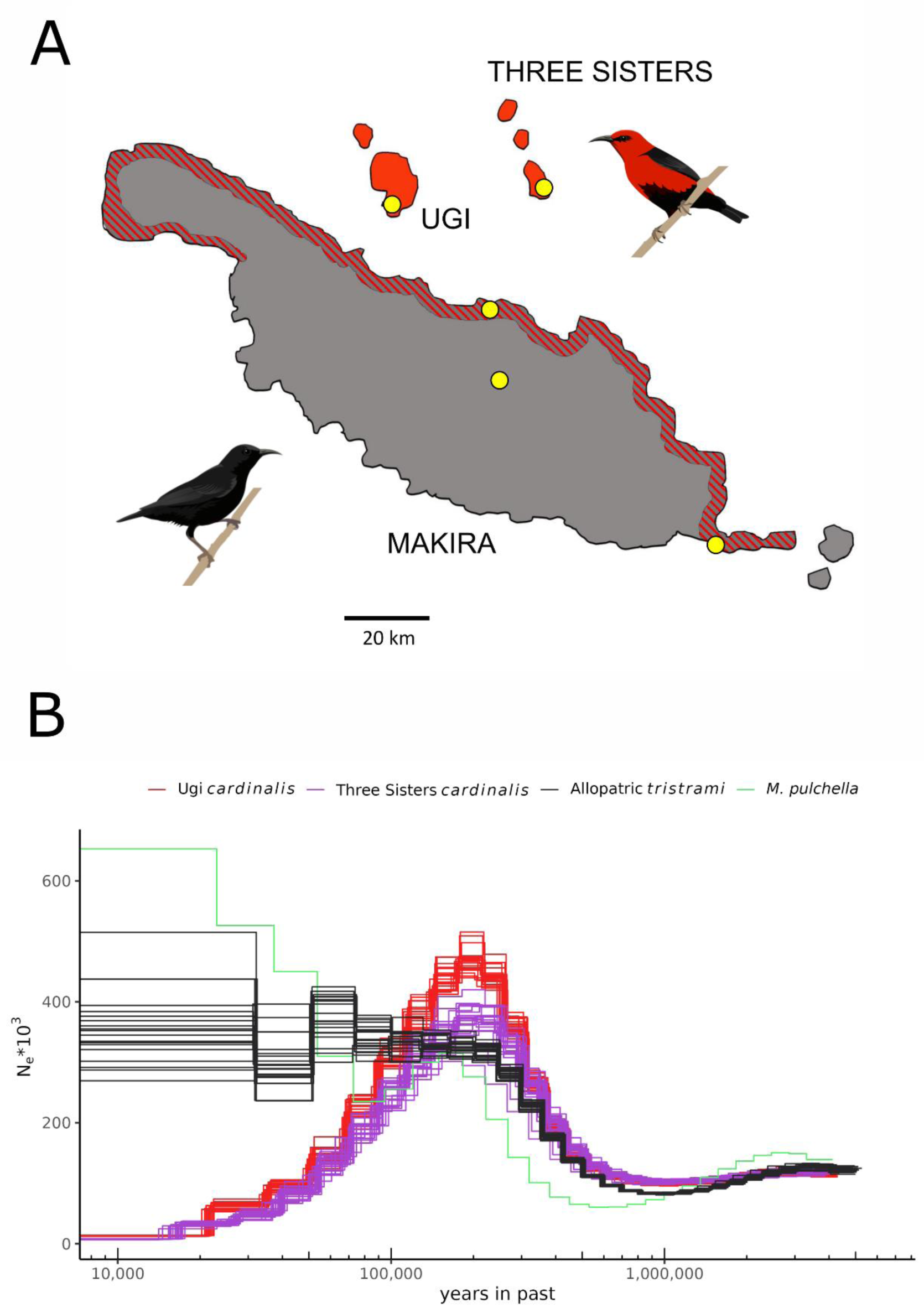
(A) Sampling sites (yellow dots) for allopatric *M. tristrami* (black bird, gray regions), allopatric *M. cardinalis* (red bird, red regions), and sympatric individuals of both species and phenotypic hybrids (red and gray striped regions). (B) PSMC inferred demographic history using autosomes of allopatric individuals show similar, declining effective populations sizes for *M. cardinalis* on Ugi and Three Sisters, while *M. tristrami* and the outgroup species *M. pulchella* have maintained or increased effective population size in recent past.

Leveraging whole-genome resequence data, we revisit this classic case of recent secondary contact and address three questions. (1) What is the history of secondary contact between *M. cardinalis* and *M. tristrami*? (2) What is the extent of admixture in the newly established region of sympatry? and (3) How does the amount and direction of introgression vary across the genome and, in particular, on sex and neo-sex chromosomes? To answer these questions, we use a newly assembled reference genome for *M. tristrami*, and whole-genome resequencing for 143 individuals, including samples from both allopatric and sympatric populations of each species. We find evidence for bidirectional introgression of autosomal loci, limited introgression of ancestral Z and especially neo-Z sequence, and asymmetric introgression of ancestral W and neo-Ws from *M. cardinalis* into *M. tristrami.* Although gene flow occurs, limited and asymmetric introgression of sex and neo-sex chromosomes highlight the potential for these genomic regions to maintain species boundaries in sympatry.

## Results and Discussion

### Neo-sex chromosomes formed by fusion of sex chromosomes with chromosome 5

We generated a highly contiguous, chromosome-level reference assembly for *Myzomela tristrami* (*Mtris*) based on PacBio HiFi long-reads. The primary assembly had an N50 of 25.7Mb and a total length of 1505.7Mb (S1 Table). After scaffolding and removal of autosomal haplotigs, we conducted a quantitative assessment of conserved avian single-copy orthologs using BUSCO [65], finding an overall BUSCO completeness score of 96.6%. The completed reference genome comprises 44 chromosomes and has a length of 1257.8Mb, including both sex chromosomes. We detected two chromosome fusion events at the sex chromosomes. The Z and W chromosomes are both fused to approximately 86% of chromosome 5, while the remaining approximately 14% assembles separately and apparently continues as autosomal sequence (chr5_remnant). Two-thirds of the now W-linked former chromosome 5 segment is distinguished by multiple rearrangements and extensive sequence divergence from its Z-linked counterpart, indicating that this region no longer recombines. We therefore refer to these segments as neo-Z and neo-W regions, respectively. Recombination between the *Mtris* sex chromosomes is mediated by a new pseudo-autosomal region derived from the remainder of the newly added chromosome 5 sequence and is referred to here as the neo-PAR. Detailed analyses of the origins, evolution, and structure of the neo-sex chromosomes in these species will be reported elsewhere.

### Allopatric populations show distinct demographic histories

For population genomics analyses, we generated short-read sequence data for a total of 143 individuals: 60 sampled from allopatric and 70 from sympatric regions of the *Mtris* and *M. cardinalis* (*Mcard*) ranges; 12 phenotypic hybrids; and 1 individual from an outgroup species, *M. pulchella* (*Mpulc*; S2 Table; Fig 1A). After mapping and filtering for quality and depth (see Methods), our dataset consisted of 30,283,937 single nucleotide polymorphisms (S3 Table).

To infer the speciation and demographic histories of the allopatric populations of *Mtris* (Highland) and *Mcard* (Ugi and Three Sisters) and the outgroup *Mpulc,* we used pairwise sequential Markov coalescent (PSMC) analyses of the autosomes [66]. These analyses suggest that after the three species diverged, they experienced a period of sustained expansion (Fig 1B). While the inferred effective population size (*N*_e_) of *Mtris* has been relatively stable over the past ∼100 Ky, the inferred *N*_e_ for *Mcard* shows evidence of steady decline. The widespread distribution of *Mcard* subspecies across south Pacific islands suggests Ugi and Three Sisters populations are at the leading edge of the species’ range [4]. We therefore infer that the reduction in *N*_e_ reflects its history of serial founder events during island colonization.

The smaller *N*_e_ of the allopatric *Mcard* versus *Mtris* is consistent with the lower average nucleotide diversities across all genomic compartments, including autosomes, neo-PAR, Z, neo-Z, W, neo-W, and mitochondria (Table 1, S4 Table). Tajima’s *D* for *Mtris* autosomes show an excess of rare SNPs consistent with modest recent population growth, whereas *Mcard* autosomes show an excess of intermediate-frequency SNPs consistent with a recent reduction in *N*_e_ (Table 2). Sex chromosome diversity is considerably lower than autosomal diversity in both species. The Z/A ratio, for example, is much lower than the 0.75 expected on the assumptions of equal sex ratios, random mating, and equal mutation rates in the two sexes: for allopatric *Mtris*, Z/A = 0.646, whereas for Ugi *Mcard*, Z/A = 0.452 and for Three Sisters *Mcard*, Z/A = 0.298 (for autosomes in these ratios, we used only chromosomes 1-10; S5 Table; see Methods). The lower Z/A ratio in allopatric *Mcard* is also consistent with a population bottleneck in its recent history [67]. Together, these data suggest that the allopatric *Mcard* populations are relatively recent arrivals to Ugi and the Three Sisters. Their most recent dispersal event was to Makira, where they encountered an historically stable population of the endemic *Mtris*. This is consistent with the phenotypic observations of expeditions in the 20^th^ century and the hypothesized history proposed in the literature [4]. We next turn to the genomic consequences of secondary contact.

**Table 1:** Nucleotide diversity, *π*.

| population | n | autosome | neo-PAR | Z | neo-Z | W | neo-W | mtDNA |
| --- | --- | --- | --- | --- | --- | --- | --- | --- |
| *Myzomela cardinalis* |  |  |  |  |  |  |  |  |
| Allopatry | 40 | 0.0023<br>(7e-06) | 0.0025<br>(3.8e-05) | 0.0009<br>(1.9e-05) | 0.0008<br>(2e-05) | 5e-06<br>(1e-06) | 4e-06<br>(0) | 0.0004<br>(NA) |
| Sympatry | 40 | 0.0025<br>(7e-06) | 0.0026<br>(4.1e-05) | 0.0010<br>(1.9e-05) | 0.0007<br>(2.1e-05) | 5e-06<br>(1e-06) | 4e-06<br>(0) | 0.0002<br>(NA) |
| *Myzomela tristrami* |  |  |  |  |  |  |  |  |
| Allopatry | 20 | 0.0028<br>(9e-06) | 0.0034<br>(4.2e-05) | 0.0017<br>(2e-05) | 0.0015<br>(1.7e-05) | 2.3e-05<br>(2e-06) | 2e-05<br>(0) | 0.0013<br>(NA) |
| Sympatry | 30 | 0.0029<br>(9e-06) | 0.0034<br>(4.1e-05) | 0.0018<br>(2.1e-05) | 0.0015<br>(1.7e-05) | 0.0006<br>(1.5e-05) | 0.0006<br>(4e-06) | 0.0138<br>(NA) |
| Nucleotide diversity ($\pi$) averaged across 50 kb windows, standard error in parentheses for each sampled population of phenotypic parental *Myzomela cardinalis* and *M. tristrami*. Number of individuals for each species/sampling region shown in column n. | | | | | | | | |

**Table 2:**
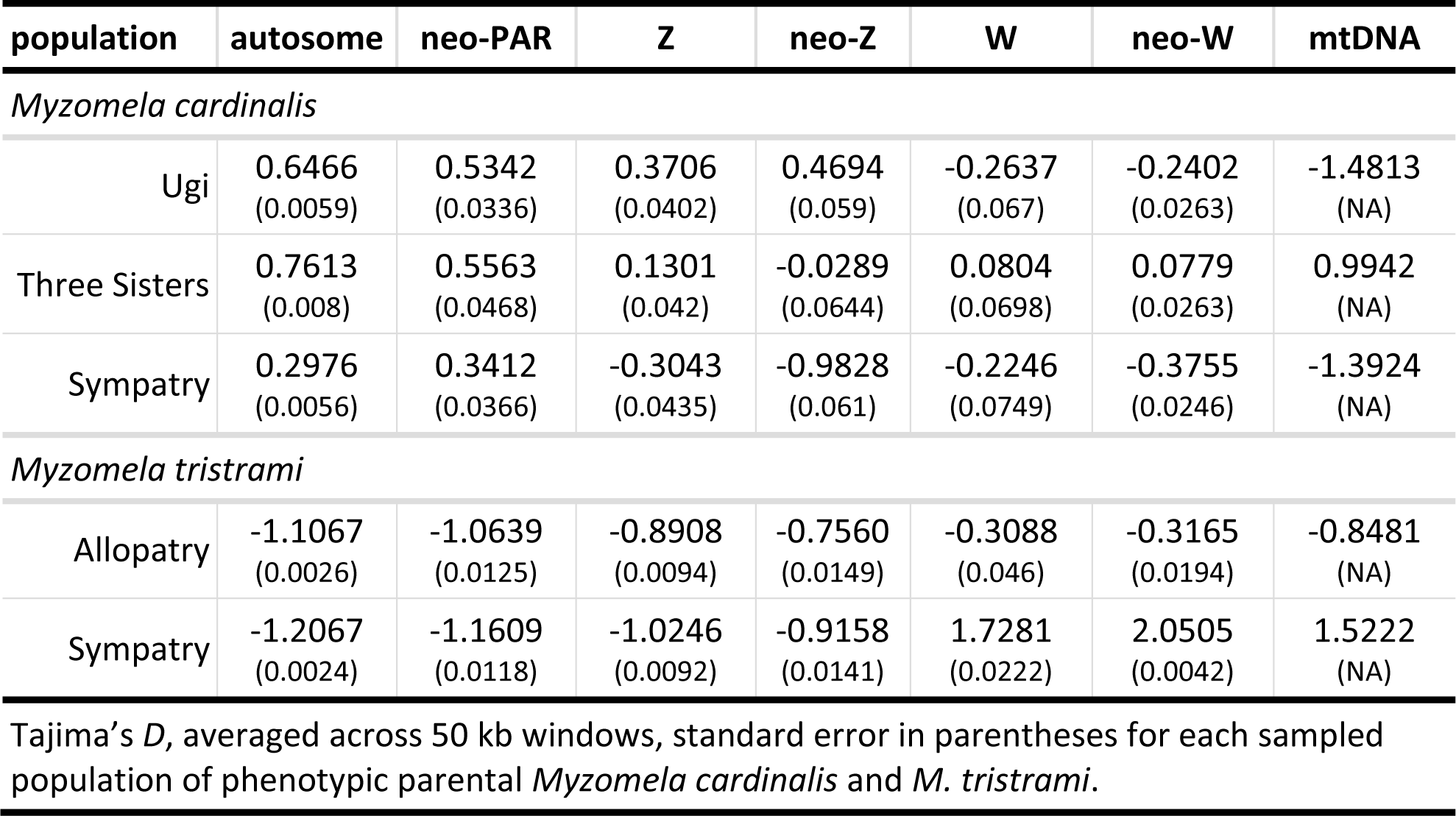
Tajima’s *D*.

| population | autosome | neo-PAR | Z | neo-Z | W | neo-W | mtDNA |
| --- | --- | --- | --- | --- | --- | --- | --- |
| *Myzomela cardinalis* |  |  |  |  |  |  |  |
| Ugi | 0.6466<br>(0.0059) | 0.5342<br>(0.0336) | 0.3706<br>(0.0402) | 0.4694<br>(0.059) | -0.2637<br>(0.067) | -0.2402<br>(0.0263) | -1.4813<br>(NA) |
| Three Sisters | 0.7613<br>(0.008) | 0.5563<br>(0.0468) | 0.1301<br>(0.042) | -0.0289<br>(0.0644) | 0.0804<br>(0.0698) | 0.0779<br>(0.0263) | 0.9942<br>(NA) |
| Sympatry | 0.2976<br>(0.0056) | 0.3412<br>(0.0366) | -0.3043<br>(0.0435) | -0.9828<br>(0.061) | -0.2246<br>(0.0749) | -0.3755<br>(0.0246) | -1.3924<br>(NA) |
| *Myzomela tristrami* |  |  |  |  |  |  |  |
| Allopatry | -1.1067<br>(0.0026) | -1.0639<br>(0.0125) | -0.8908<br>(0.0094) | -0.7560<br>(0.0149) | -0.3088<br>(0.046) | -0.3165<br>(0.0194) | -0.8481<br>(NA) |
| Sympatry | -1.2067<br>(0.0024) | -1.1609<br>(0.0118) | -1.0246<br>(0.0092) | -0.9158<br>(0.0141) | 1.7281<br>(0.0222) | 2.0505<br>(0.0042) | 1.5222<br>(NA) |
| Tajima's *D*, averaged across 50 kb windows, standard error in parentheses for each sampled population of phenotypic parental *Myzomela cardinalis* and *M. tristrami*. |  |  |  |  |  |  |  |

### Autosomal loci introgress in both directions at secondary contact

We captured 187 birds in sympatry. Of these, 68 were phenotypically *Mtris*, 107 were phenotypically *Mcard*, and 12 were identified as “phenotypic hybrids” — individuals with plumage characteristics clearly intermediate between *Mtris* and *Mcard* (mostly black with some red feathers). These individuals raise the possibility that our sample includes backcross or advanced generation hybrids that are indistinguishable from the parental species. To test for the possibility of cryptic hybrids, we start by focusing on analyses of autosomal regions of the genome.

We used five approaches to characterize autosomal admixture. First, simple summaries of differentiation and divergence restricted to phenotypically parental individuals (excluding phenotypic hybrids) support admixture between sympatric populations (Table 3, S8, S9 Table). Autosomal differentiation (*F*_ST_) between allopatric *Mcard* and *Mtris* is 0.282, whereas that for sympatric *Mcard* and *Mtris* drops to *F*_ST_ = 0.197 (Table 3). Absolute divergence between species changes less dramatically in sympatry but does shift from *d*_xy_ ≈ 0.0035 in allopatry to *d*_xy_ = 0.0033 in sympatry (S8 Table). These observations are consistent with introgression between the two species in sympatry reducing differentiation and divergence even when looking solely at birds with purely parental phenotypes.

**Table 3:** Population differentiation, *F*_ST_.

| population comparison | autosome | neo-PAR | Z | neo-Z | W | neo-W | mtDNA |
| --- | --- | --- | --- | --- | --- | --- | --- |
| *Myzomela cardinalis* |  |  |  |  |  |  |  |
| Allopatry vs. Sympatry | 0.0323<br>(2e-04) | 0.0326<br>(0.0013) | 0.0508<br>(0.0016) | 0.0668<br>(0.0033) | -0.0067<br>(0.0061) | -0.0032<br>(0.002) | 0.0019<br>(NA) |
| *Myzomela tristrami* |  |  |  |  |  |  |  |
| Allopatry vs. Sympatry | 0.0061<br>(1e-04) | 0.0047<br>(2e-04) | 0.0054<br>(3e-04) | 0.0051<br>(5e-04) | 0.1548<br>(8e-04) | 0.1557<br>(2e-04) | 0.1233<br>(NA) |
| Heterospecific |  |  |  |  |  |  |  |
| Allo. *cardinalis* vs. Allo. *tristrami* | 0.2818<br>(0.001) | 0.2757<br>(0.0061) | 0.5921<br>(0.0048) | 0.6606<br>(0.0052) | 0.9917<br>(7e-04) | 0.9924<br>(2e-04) | 0.9831<br>(NA) |
| Allo. *cardinalis* vs. Sym. *tristrami* | 0.2423<br>(9e-04) | 0.2426<br>(0.0054) | 0.5704<br>(0.0048) | 0.644<br>(0.0049) | 0.7409<br>(8e-04) | 0.7411<br>(2e-04) | 0.8017<br>(NA) |
| Allo. *tristrami* vs. Sym. *cardinalis* | 0.2354<br>(9e-04) | 0.2509<br>(0.0066) | 0.5717<br>(0.005) | 0.6630<br>(0.0067) | 0.9920<br>(6e-04) | 0.9927<br>(2e-04) | 0.9855<br>(NA) |
| Sym. *cardinalis* vs. Sym. *tristrami* | 0.1972<br>(8e-04) | 0.2178<br>(0.0059) | 0.5511<br>(0.005) | 0.6433<br>(0.0064) | 0.7411<br>(7e-04) | 0.7413<br>(2e-04) | 0.8041<br>(NA) |

$F_{ST}$ averaged across 50kb windows for each region of the genome, for pairwise comparisons of phenotypic parental populations of *Myzomela cardinalis* and *M. tristrami*. Allopatry and sympatry abbreviated as Allo. and Sym., respectively. Standard errors in parentheses.

Second, to infer the amount and direction of introgression between species we conducted ABBA-BABA analyses using the program Dsuite [68] to calculate the *D-*statistic and *f_4_* admixture ratio (Table 4). ABBA-BABA uses counts of the distribution of derived alleles shared among four taxa using the topology (((P1,P2)P3)P4) to determine if gene flow has occurred between the three ingroup taxa [69]. We used *Mpulc* as the P4 outgroup and analyzed two different topologies: “*cardinalis* P3” with allopatric *tristrami* (P1), sympatric *tristrami* (P2), and sympatric *cardinalis* (P3) as the ingroups; and *“tristrami* P3” with allopatric *cardinalis* (P1), sympatric *cardinalis* (P2), and sympatric *tristrami* (P3) as the ingroups. For both topologies, *D* statistics were positive and significant, indicating excess sharing of derived alleles between sympatric *cardinalis* and *tristrami* consistent with autosomal gene flow in both directions. A slightly higher admixture proportion (*f_4_* ratio) calculated for the *tristrami* P3 topology (0.15) indicates more gene flow from sympatric *Mtris* into sympatric *Mcard* than the reverse (0.09; Table 4). Further parsing signals of introgression by chromosome shows that *D* statistics are significant for both topologies for most autosomes (74% for *tristrami* P3, 95% for *cardinalis* P3; Fig 2, S11 Figure).

**Figure 2.**
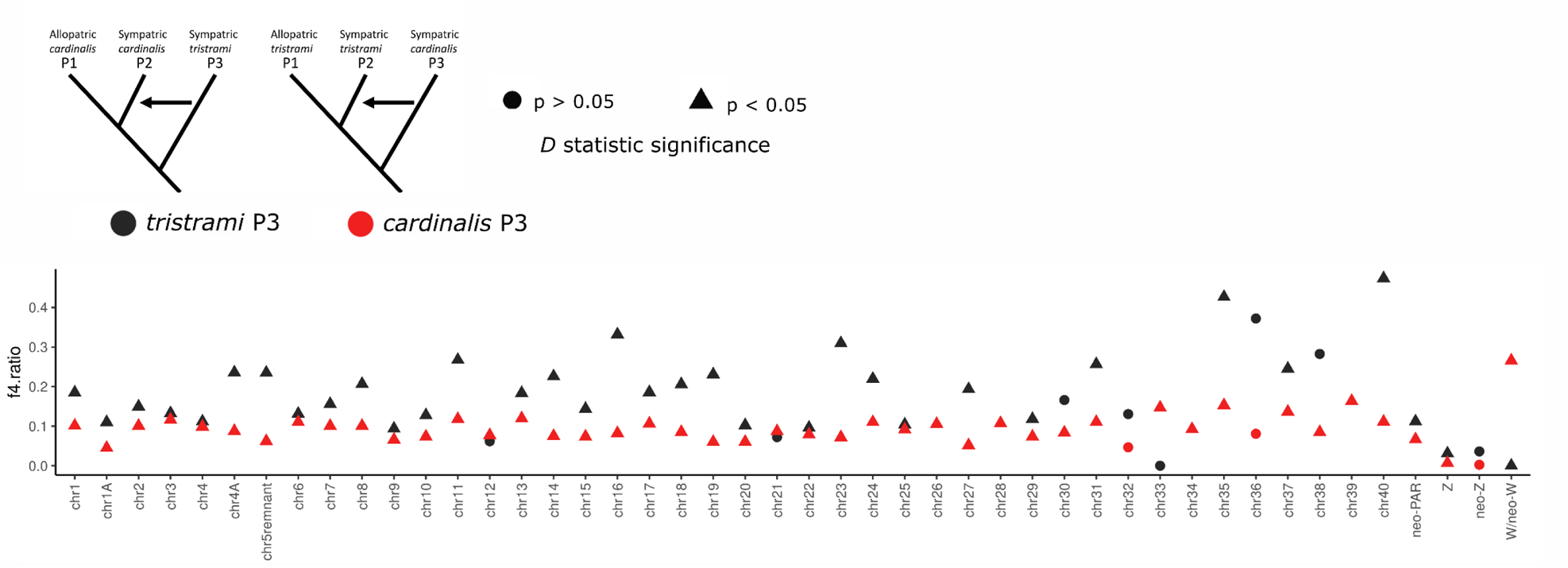
Admixture ratio (*f_4_* statistic) for each autosome and sex chromosome regions. Color of the point indicates which topology the statistic was calculated for (*tristrami* P3 or *cardinalis* P3), and shape of the point indicates whether the *D* statistic for that chromosome was significantly different from zero, using the block-jackknife procedure. Admixture ratios calculated for *tristrami* P3 taxa on chr 26, 28, 34, and 39 used an alternative topology indicating nonsignificant introgression from sympatric *tristrami* into allopatric *cardinalis,* shown in S11 Figure.

**Table 4:**
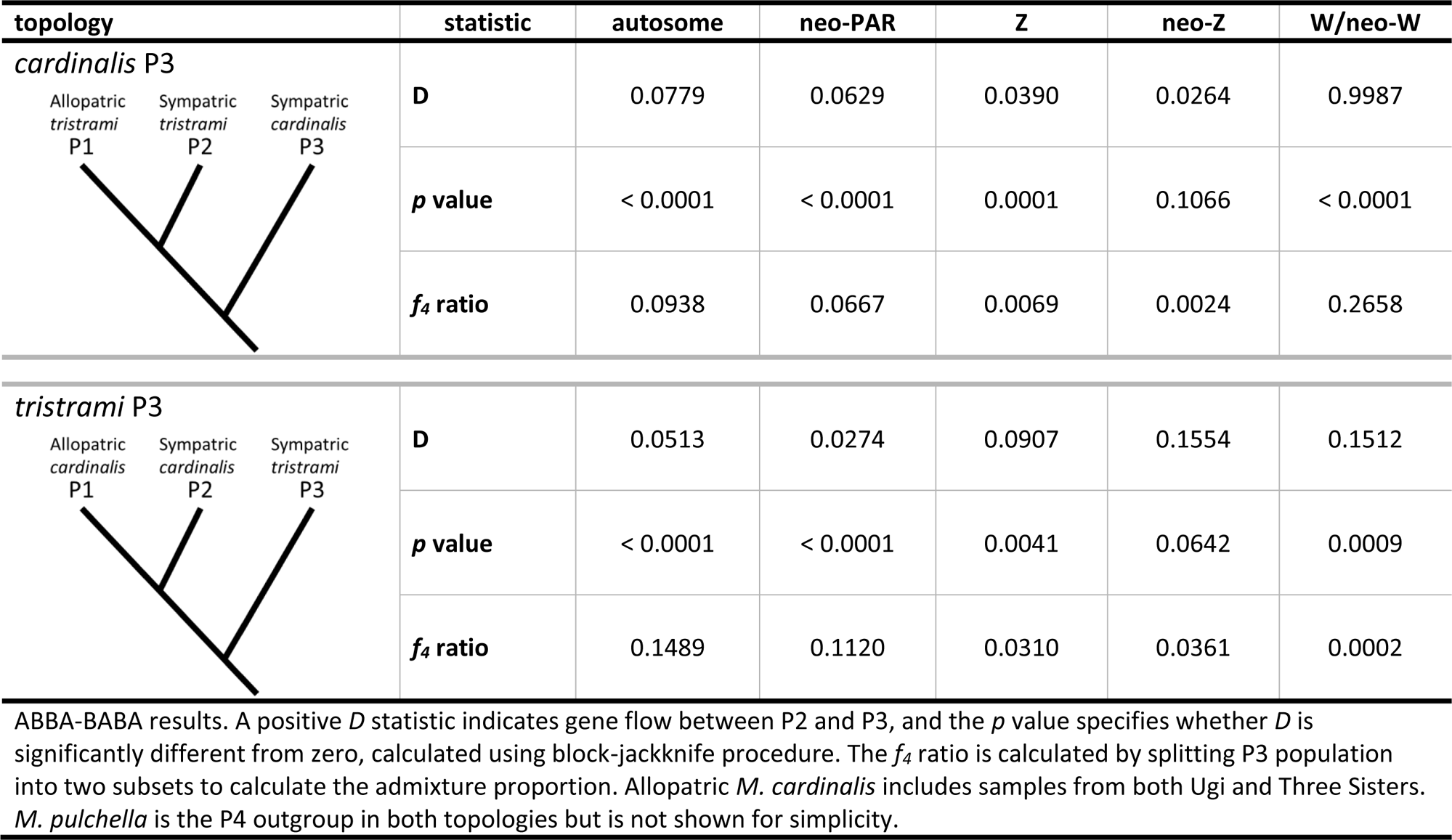
Test of introgression, ABBA-BABA *D-*statistics.

| topology | statistic | autosome | neo-PAR | Z | neo-Z | W/neo-W |
| --- | --- | --- | --- | --- | --- | --- |
| *cardinalis* P3 | **D** | 0.0779 | 0.0629 | 0.0390 | 0.0264 | 0.9987 |
|  | ***p* value** | < 0.0001 | < 0.0001 | 0.0001 | 0.1066 | < 0.0001 |
|  | ***f*<sub>4</sub> ratio** | 0.0938 | 0.0667 | 0.0069 | 0.0024 | 0.2658 |
| *tristrami* P3 | **D** | 0.0513 | 0.0274 | 0.0907 | 0.1554 | 0.1512 |
|  | ***p* value** | < 0.0001 | < 0.0001 | 0.0041 | 0.0642 | 0.0009 |
|  | ***f*<sub>4</sub> ratio** | 0.1489 | 0.1120 | 0.0310 | 0.0361 | 0.0002 |

ABBA-BABA results. A positive *D* statistic indicates gene flow between P2 and P3, and the *p* value specifies whether *D* is significantly different from zero, calculated using block-jackknife procedure. The *f*<sub>4</sub> ratio is calculated by splitting P3 population into two subsets to calculate the admixture proportion. Allopatric *M. cardinalis* includes samples from both Ugi and Three Sisters. *M. pulchella* is the P4 outgroup in both topologies but is not shown for simplicity.

Third, we used principal component analyses (PCA) to identify admixed individuals. PCA revealed clear separation of *Mcard* and *Mtris* species along principal component 1 (PC1). As expected, phenotypic hybrids were intermediate in PC1 values, but several phenotypic *Mcard* and *Mtris* individuals also fell between the two clusters of parental species and are likely backcross or advanced generation hybrids (Fig 3A).

**Figure 3.**
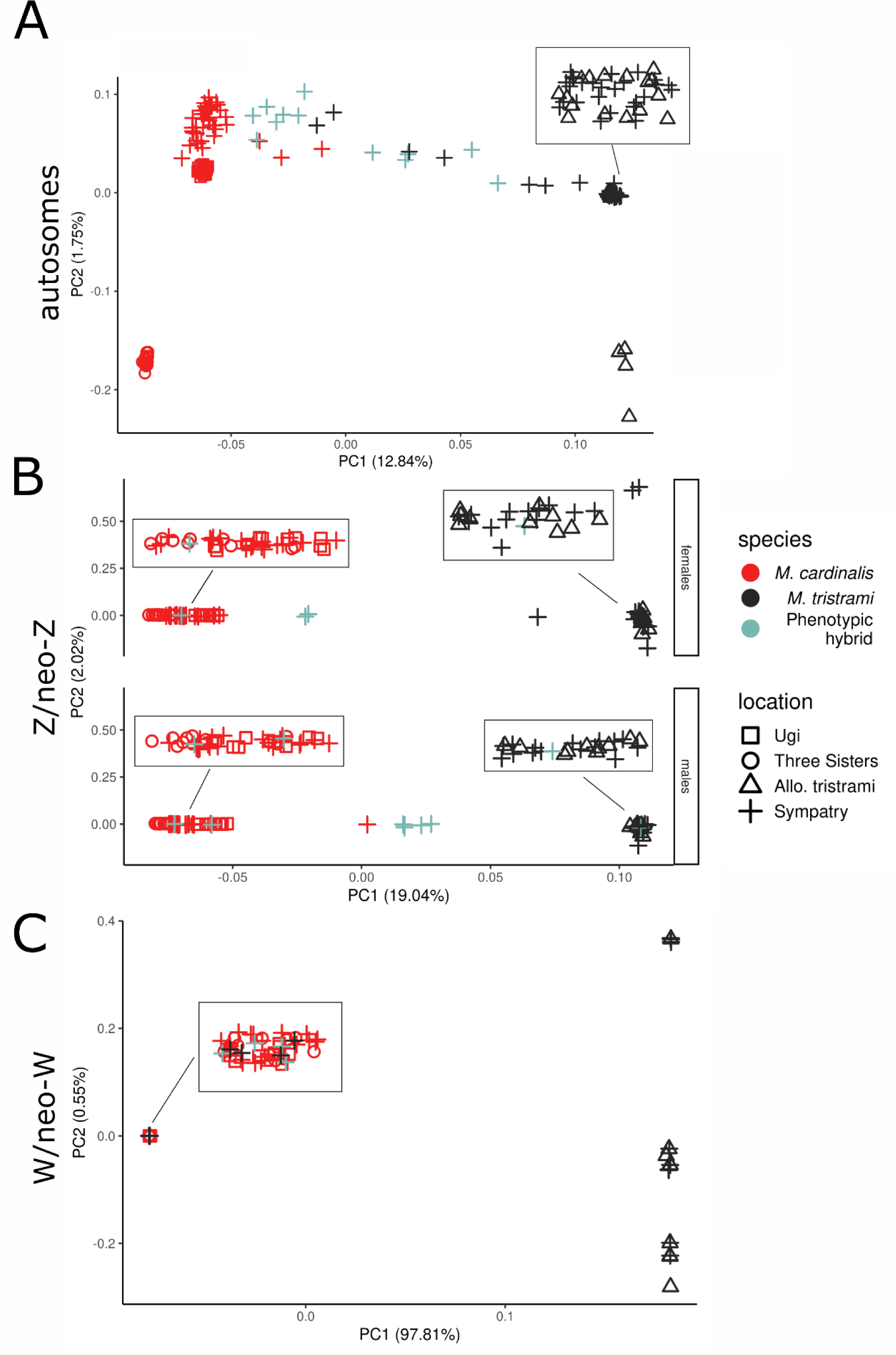
Principal components analysis of autosomal (A), Z and neo-Z (B) and W and neo-W (C) sequence. Symbol color represents phenotypic species assignment while symbol shape indicates sampling locality. Inset boxes show points jittered for visualization. Plot for Z/neo-Z is separated by sex to distinguish homogametic males and heterogametic females.

To explicitly test for production of F_1_s and advanced generation backcrosses we used SNPs fixed between allopatric *Mcard* and *Mtris* to calculate interspecific heterozygosity and hybrid index for sympatric individuals. We see clear evidence of F_1_s with maximized interspecific heterozygosity and hybrid index ≈ 0.5. The majority are phenotypic hybrids (n = 8), but phenotypic *Mcard* (n = 2) and *Mtris* (n = 1) also appear to be F_1_ individuals. There is also evidence of backcrossing to both parental species (Fig 4A).

**Figure 4.**
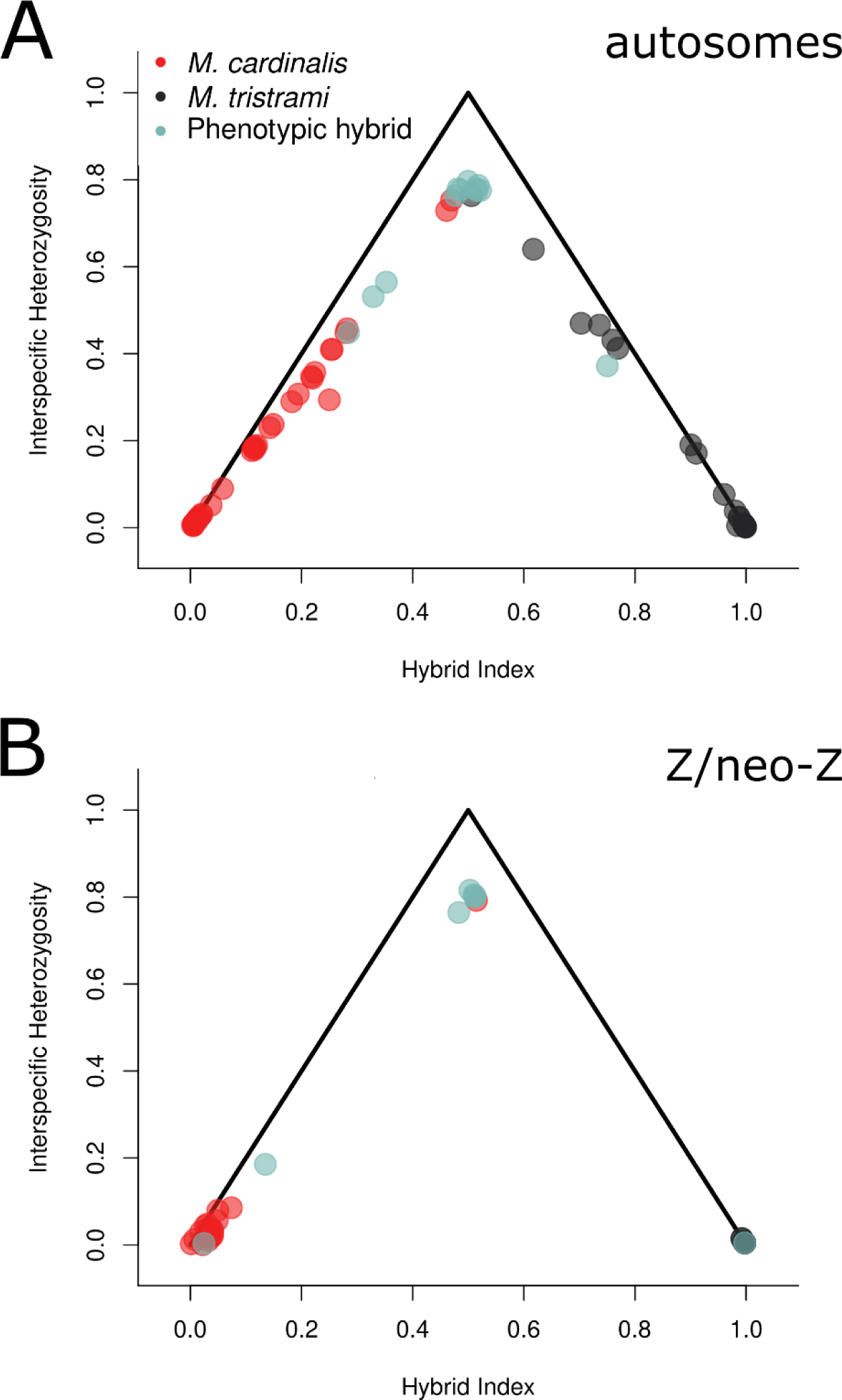
Triangle plots showing the relationship between interspecific heterozygosity and hybrid index calculated using 2449 autosomal (A) and 37,613 Z/neo-Z (B) SNPs fixed between species in allopatry. Circles are sympatric individuals of both sexes (A) or sympatric males (B), colored by phenotype (see legend). F1 individuals fall at the maximum heterozygosity and hybrid index ≈ 0.5, while advanced generation backcrosses fall along the legs of the triangle as interspecific heterozygosity declines and hybrid index is closer to 0 (*M. cardinalis* ancestry) or increases to 1 *(M. tristrami* ancestry).

Finally, we used ADMIXTURE to estimate individual ancestry proportions [70]. Cross-validation error was minimized when the number of groups (*K*) was equal to two (S10 Figure). However, *K* = 3 was also well supported for autosomal sequence and provided insight into structuring between Ugi and Three Sisters *Mcard* (Fig 5A, see below for further discussion of *Mcard* allopatric populations). All phenotypic hybrids, as well as several individuals in sympatry (both phenotypically *Mtris* and phenotypically *Mcard*) show autosomal ancestry from both species. In summary, autosomal sequence indicates extensive admixture between *Mcard* and *Mtris* in sympatry. Advanced generation hybrids in our sample confirm that some F_1_ hybrids are fertile. The relative abundance of *Mcard* in sympatry (107 of 187 captured individuals) and continued production of phenotypic hybrids and admixed birds suggests ecological incompatibilities are not strongly influencing reproductive isolation in the region of sympatry, although additional work is necessary to confirm this.

**Figure 5.**
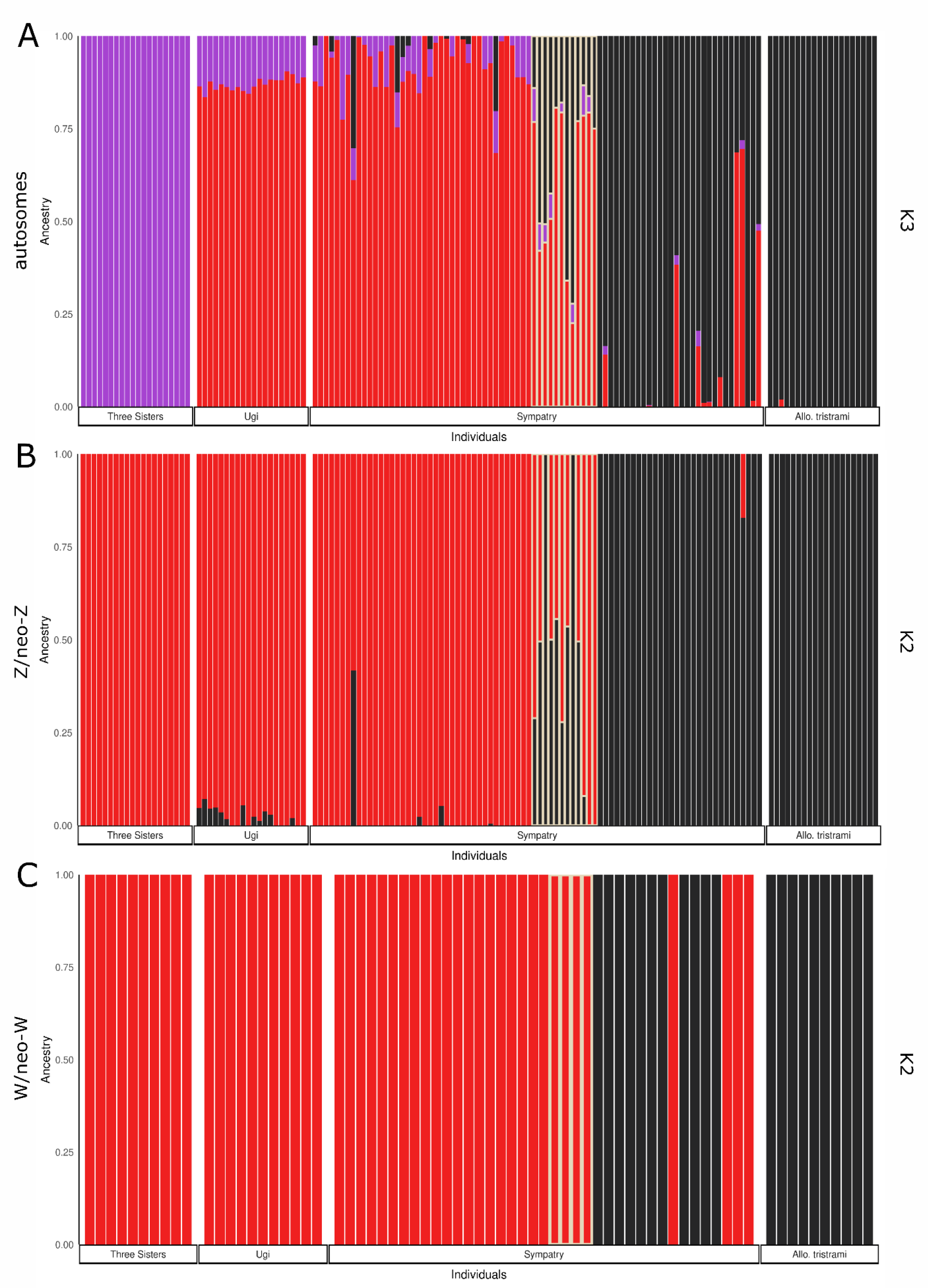
Proportion ancestry calculated in ADMIXTURE for autosomes (A), Z/neo-Z (B), and W/neo-W (C). Autosomes are shown at K = 3 (see S9 Figure for K = 2), while Z/neo-Z and W/neo-W shown at K = 2. Individuals are grouped by sampling location. Phenotypic hybrids are outlined in yellow, with phenotypic *M. cardinalis* to the left and phenotypic *M. tristrami* to the right of phenotypic hybrids.

### New pseudo-autosomal region shares patterns of introgression with autosomes

The new pseudo-autosomal region, or neo-PAR, is a region of the former autosome that is now linked to the sex chromosomes but has continued to recombine in both sexes. As such, it serves as an important point of comparison with neo-sex chromosome regions that have ceased to recombine. The neo-PAR has nucleotide diversity similar to autosomes (Table 1). Differentiation within and between species on the neo-PAR is intermediate to autosomes and other regions of the sex chromosomes (Table 3). Finally, signals of introgression on the neo-PAR echo those of the autosomes, with significant *D* statistics for both topologies and admixture ratios that were only slightly lower for the neo-PAR than for autosomes, indicating bidirectional gene flow (Table 4). Thus, despite transitioning from autosome to sex chromosome, this region appears to continue to share characteristics with autosomes and shows little evidence of being a barrier to introgression. We now compare the pattern of bidirectional autosomal and neo-PAR gene flow to introgression patterns observed on the sex and neo-sex chromosomes.

### Z /neo-Z chromosome are refractory to introgression

The Z/neo-Z chromosome shows evidence of admixture, but the degree of introgression is markedly reduced compared to autosomes. Differentiation on the Z and neo-Z between *Mcard* and *Mtris* is slightly lower in sympatry (Z: *F*_ST_ = 0.551, neo-Z: *F*_ST_ = 0.643) than in allopatry (Z: *F*_ST_ = 0.592, neo-Z: *F*_ST_ = 0.661), but overall much higher than autosomes (Table 3). Divergence between *Mcard* and *Mtris* in allopatry is similar to that in sympatry. In the ABBA-BABA analysis, *D* statistics for ancestral Z sequence were significant, consistent with gene flow between sympatric *Mcard* and *Mtris*, but *f_4_* admixture ratios were much lower than those estimated for autosomal sequence (Table 4, Fig 2). By contrast, *D* statistics for the neo-Z provide no evidence for gene flow.

Analyses of individual genotypes revealed F_1_, backcross, and advanced generation hybrids in our sample. For PCA of the Z/neo-Z, we separated homogametic males and heterogametic females for visualization (Fig 3B). *Mcard* and *Mtris* clearly separated along PC1. Five of eight phenotypic hybrid males and one phenotypic *Mcard* male were intermediate in PC1, potentially F_1_ individuals heterozygous for species’ Z/neo-Z haplotypes. However, there were also phenotypic hybrid males which fell within the species’ clusters at either end of PC1, indicating those individuals may be advanced generation hybrids homozygous for Z/neo-Z haplotypes from either *Mcard* (n = 2) or *Mtris* (n = 1). Although females only have one Z haplotype, we do see individuals falling toward the center of PC1 (Fig 3B). These females are likely backcross individuals carrying a recombinant Z/neo-Z containing both *Mcard* and *Mtris* sequence within a single haplotype. Plotting hybrid index against interspecific heterozygosity for diploid males confirmed the majority of phenotypic hybrid males and one phenotypic *Mcard* male were heterozygous for Z/neo-Z haplotype (Figure 4B). Individual-level estimates of ancestry proportions of Z/neo-Z calculated in ADMIXTURE showed some but not all phenotypic hybrids to be admixed, while only two phenotypically parental individuals had appreciable ancestry from the other species (Fig 5B).

Together, these results show introgression of Z/neo-Z in sympatry is limited relative to autosomal sequence. Aggregated, population-level estimates of differentiation are similar between allopatric and sympatric species comparisons, while estimates of introgression and admixture proportion are very low, and in the case of the neo-Z, not significant. At the individual level, only a few phenotypic hybrid individuals carry genetic material from both species, suggesting this region of the genome is refractory to gene flow between *Mcard* and *Mtris*.

### Strong, asymmetric introgression of W/neo-W and mitochondria

Genomic sequence for the W, neo-W, and the mitochondria are maternally co-inherited and the two species carry distinct sets of haplotypes. All three regions show strong, asymmetric introgression from *Mcard* into *Mtris.* Interspecific differentiation between *Mtris* and *Mcard* is lower in sympatry (*F*_ST_ = 0.74) than in allopatry for W and neo-W (*F*_ST_ = 0.99; Table 3), consistent with introgression reducing differentiation between species where gene flow is possible. Comparing intraspecific differentiation across genomic compartments, *Mtris* shows much higher differentiation for W/neo-W (*F*_ST_ = 0.15) and mitochondrial sequence (*F*_ST_ = 0.12) relative to autosomal sequence (*F*_ST_ < 0.01; Table 3). These findings imply the presence of foreign *Mcard* W/neo-W and mitochondrial haplotypes in sympatric *Mtris* females. ABBA-BABA analyses for W/neo-W haplotypes further support asymmetric introgression from *Mcard* into *Mtris.* The *cardinalis* P3 topology showed a significant *D* statistic and an *f_4_* admixture proportion of 0.27; *tristrami* P3 topology also had a significant *D* statistic, but admixture proportion was ≈ 0 (Table 4, Fig 2). Movement of *Mcard* W/neo-W haplotypes into *Mtris* is also evident from analyses of individual genotypes. The PCA plots reveal that all phenotypic hybrid females and four of fifteen phenotypic *Mtris* females cluster with the *Mcard* individuals, whereas none of the sympatric *Mcard* females cluster with *Mtris* (Fig 3C). Therefore, none of the phenotypic *Mtris* females carry *Mcard* W/neo-W haplotypes. The mitochondrial haplotype network, as expected, echoes the findings from the W/neo-W sequence: haplotypes for *Mtris* are restricted to phenotypic *Mtris,* whereas *Mcard* mitochondrial haplotypes are carried by phenotypic hybrids and sympatric *Mtris* (Fig 6). These results are consistent with previous analyses of mitochondrial markers [61]. Finally in the ADMIXTURE analyses, all sympatric *Mcard,* phenotypic hybrids, and four of the fifteen sympatric *Mtris* females (∼26%) carry W/neo-W chromosomes of *Mcard* and none of the *Mcard* females show any *Mtris* ancestry (Fig 5C).

**Figure 6.**
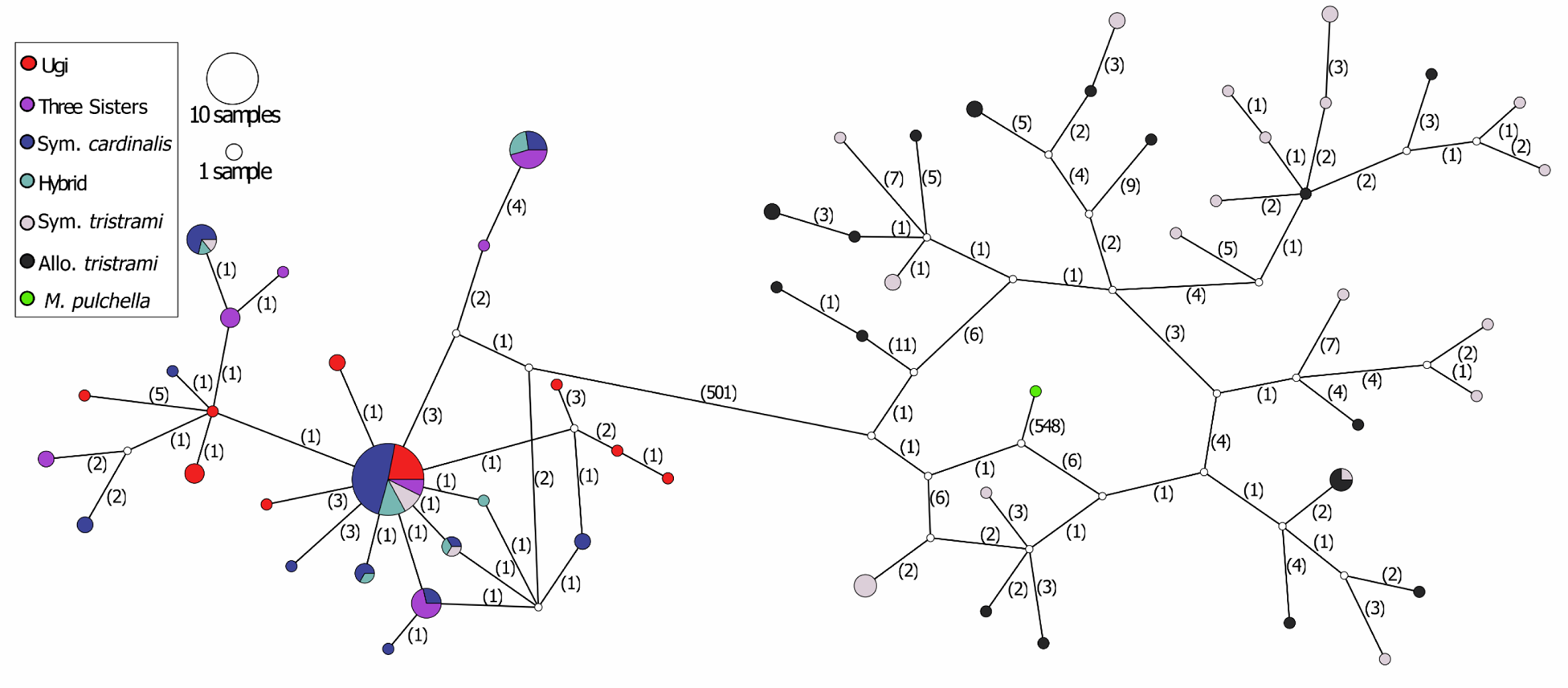
Mitochondrial haplotype TCS network show that *M. cardinalis* haplotypes are much less diverse than *M. tristrami* haplotypes but shared by hybrids and phenotypic *M. tristrami* individuals in sympatry (mutations between nodes shown in parentheses).

Consistent with asymmetric introgression, our sample has four phenotypic hybrid females and four phenotypic *Mtris* females produced by crosses between *Mcard* females and *Mtris* males, but none from the reciprocal cross. The absence of hybrid or *Mcard* females with *Mtris* W/neo-W or mtDNA in our dataset, and hence the apparent lack of introgression in that direction has two potential explanations. First, if matings between *Mtris* females and *Mcard* males occur, Haldane’s rule for lethality predicts a dearth of F_1_ hybrid daughters [38]; in addition, accumulation of Dobhzansky-Muller incompatibilities can lead to asymmetry in such inviability [71]. However, there are several reasons to consider alternative explanations. For one, Haldane’s rule for hybrid sterility in both directions tends to evolve before hybrid lethality [72]. In addition, hybrid lethality in birds tends to occur in much older species pairs [3,73]. It remains possible, however, that the rapid evolution of the neo-W has accelerated the evolution of hybrid lethality [46].

The second explanation for asymmetric introgression is sexual incompatibilities — asymmetry in mate choice or availability which limits crosses between *Mtris* females and *Mcard* males [37]. Mayr and Diamond [4] proposed that hybridization initially occurred in sympatry due to a lack of conspecific mates for recently arrived *Mcard*. However, the relative frequency of *Mcard* in sympatry has surpassed that of the native *Mtris,* so it seems unlikely that *Mcard* females are constrained to choose heterospecific males to pair with [61]. Nevertheless, phenotypic hybrids and admixed individuals continue to be observed decades after colonization. It is therefore possible that introgression of maternally coinherited sequences from *Mcard* into *Mtris* genomic backgrounds occurs via asymmetric mate choice, is globally adaptive, or is driven by selfish W or neo-W [74,75]. Future work is necessary to distinguish these alternatives.

### More than one source for sympatric *Mcard* population

To determine if the *Mcard* population(s) that invaded Makira originated from Ugi, from the Three Sisters, or from both, we assessed relationships between the two allopatric *Mcard* and sympatric *Mcard* populations. We identified very few alleles private to sympatric *Mcard* or fixed between sympatric *Mcard* and either allopatric *Mcard* population, while many alleles were shared among all three *Mcard* populations (S6, S7 Tables). Differentiation between Ugi and Three Sisters populations of *Mcard* is higher across the genome than between either allopatric population and sympatric *Mcard* (S9 Table). For autosomal sequence, PCA shows Ugi and Three Sisters individuals separating along PC2 (Fig 3A), and ADMIXTURE analyses using *K* = 3 show distinct ancestry for Three Sisters (Fig 5A). Finally, the mitochondrial network shows that haplotypes otherwise unique to Ugi and Three Sisters are present in sympatric *Mcard,* sympatric *Mtris*, and phenotypic hybrids (Fig 6). Thus, both Ugi and Three Sisters individuals contributed to the founding population of *Mcard* on Makira.

## Conclusions

Our population genomic study of *Myzomela* honeyeaters in the Solomon Islands sheds light on a system with complex and ongoing introgression following very recent secondary contact of closely related taxa carrying neo-sex chromosomes. Sex chromosomes are known to play a large role in speciation [33]. Neo-sex chromosomes transitioning to sex-specific inheritance undergo rapid structural and molecular evolution, which may incidentally contribute to incompatibilities between species [46,47,76]. Consistent with genetic or sexual incompatibilities accumulating on sex and neo-sex chromosomes, we see limited introgression of Z/neo-Z and asymmetric introgression of W/neo-W, while autosomal regions introgress into both species. The patterns of introgression that we see highlight the potential for gene flow to vary across the genome, and the importance of sex chromosomes in maintaining species boundaries. Importantly, gene flow is especially limited for neo-Z sequence, and the strong asymmetric introgression of W/neo-W sequence may be due to rapid accumulation of incompatibilities in that region. Although further work is needed to determine the specific behaviors and/or loci that underlie these patterns, this study addresses fundamental questions in speciation research: how species persist in the face of gene flow, and what role sex and neo-sex chromosomes play in reproductive isolation. Despite hybridization, large regions of the genome can remain refractory to introgression, maintaining species boundaries. As rapidly evolving genomic regions with sex-specific inheritance, sex and neo-sex chromosomes have the potential to preserve species divergence, crucially influencing the process of speciation and the generation of biodiversity. By revisiting the *Myzomela* honeyeater system in Makira and testing predictions of the allopatric model of speciation with genomic data, we build on the work started by Mayr and Diamond [4], providing important insights on the “moment of truth” for speciation.

## Materials and methods

### Samples and sequencing

We sampled 40 allopatric *Myzomela cardinalis* from two island groups adjacent to the region of sympatry on Makira (*i.e*., Ugi and Three Sisters) and 20 allopatric *M. tristrami* from high elevation regions on the island of Makira (Fig 1A). We sampled 82 birds from the low elevation region of sympatry on Makira. Of these, 40 were identified as *M. cardinalis,* 30 were identified as *M. tristrami*, and 12 were identified as hybrids, based on a phenotype of mostly black plumage with patches of red feathers. In addition, we used a sample of *Myzomela pulchella* collected on New Ireland, Papua New Guinea and held at the University of Kansas Biodiversity Institute and Natural History Museum (see S2 Table for details on all samples).

We collected whole blood using brachial venipuncture from birds captured in mist nets at fruiting trees. We added blood to lysis buffer [77] and stored it at room temperature until arrival to the lab, where it was subsequently stored at −80 ⁰C. We extracted DNA using a Qiagen DNeasy kit with an RNase step.

Extracted genomic DNA was sequenced at Novogene (Sacramento, CA). Following quality and concentration assessment using Agarose Gel Electrophoresis and Qubit 2.0, genomic DNA was randomly fragmented and fragments were end polished, A-tailed, and ligated with Illumina adapters. Further PCR amplification preceded library construction and purification with the AMPure XP system. Finally, size distribution of libraries was checked using Agilent 2100 Bioanalyzer (Agilent Technologies, CA, USA). Libraries were then pooled and sequenced by synthesis using the Illumina platform to generate 150 bp paired end reads. The *M. pulchella* sample was sequenced at the Oklahoma Medical Research Foundation. Libraries were constructed using the Swift 2S Turbo DNA Library Kit prior to sequencing by synthesis of 150 bp paired end reads using the Illumina Novaseq machine.

### Reference genome assembly

We sequenced an *M. tristrami* female with PacBio HiFi long read and generated a *de novo* assembly using hifiasm v0.13-r308 with default parameters [78,79]. We used GeMoMa (v1.8) and the annotation from zebra finch genome bTaeGut1.4.pri to infer a rough annotation of genes in the *Myzomela* genome. We then used these rough annotations, comparing contigs against both zebra finch and the chicken genome bGalGal1.mat.broiler.GRCg7b to infer synteny relationships, remove duplicate haplotigs, and, finally, scaffold contigs into chromosomes in *Myzomela.* The resulting assembly uses the zebra finch numbering system for chromosomes 1-29; chromosome 30-40 were named in descending order of size. We generated repetitive DNA libraries using the RepeatModeler v2 pipeline [80]. RepeatModeler employs a combination of *de novo* and homology-based characterization of different classes of repeats. The repeat library was annotated and combined with Repbase, and manually curated repeat libraries from other studies [81–84].We then used RepeatMasker (Smit et al. 2013, v4.1.0) to identify repetitive regions of the genome.

### Population genomic analyses

We used Trim Galore (https://www.bioinformatics.babraham.ac.uk/projects/trim_galore/) to process raw reads. Trim Galore first removes low quality reads from the 3’ end and then trims adapter sequences using the program Cutadapt [86] before running FastQC (https://www.bioinformatics.babraham.ac.uk/projects/fastqc/) to check adapter content after trimming.

We aligned trimmed reads to the *Myzomela tristrami* reference genome using Burrows-Wheeler-Aligner (bwa-mem, v0.7.17; [87], mapping 26,893,270,155 reads and yielding a mean coverage of 16.95x. After alignment, we sorted the resulting mapped reads by coordinate using samtools, v1.7 [88]. We continued with processing following the Genome Analysis Toolkit best practices workflow (GATK 4.2.6.1 [89]). First, we used AddOrReplaceReadGroups (Picard v.2.12.0, http://broadinstitute.github.io/picard) to denote flow cell and lane of each read. We used MarkDuplicates (Picard v12.2.0) to identify duplicate reads resulting from PCR amplification. Next, we used FixMateInformation (Picard v12.2.0) to verify and correct information between mate-pairs. At this point we assessed coverage using qualimap (v2.2.1; [90]) and verified sex of individuals based on coverage of Z scaffolds (diploid in males, haploid in females). We then called variants using the GATK pipeline, starting with HaplotypeCaller, on the diploid (default) setting. Due to hemizygosity of female sex chromosomes, we first ran HaplotypeCaller on autosomes and pseudo-autosomal regions across both sexes. We then separated males and females, running HaplotypeCaller on males for Z sequence only and on females for both Z and W sequence. We then used CombineGVCFs to facilitate joint genotyping for each region of the genome. Finally, GenotypeGVCFs generated an all-sites VCF file containing both variant and invariant sites for downstream filtering and analysis.

We used VariantFiltration and GATK recommendations for hard filtering in non-model organisms to flag any sites that had QD < 2.0, SOR > 3.0, FS > 60.0, MQ < 40.0, MQRankSum < 12.5, or ReadPosRankSum < −80.0. We also flagged any sites overlapping known repetitive regions in the reference genome. We then used bcftools [88] to recode any flagged low-quality sites or sites in repetitive regions as missing. We used vcftools [91] to remove indels and assess depth of coverage prior to further filtering. We used bcftools to recode any autosomal genotypes as missing if they had less than 10x or more than 34x (twice the mean before filtering for depth) coverage averaged across all samples. For sex chromosome genotypes we adjusted our depth of coverage filters to accommodate the reduced coverage for hemizygous female samples, recoding as missing any genotypes less than 6x or more than 24x averaged across all samples. In addition, we further filtered sex chromosomes to remove any spurious heterozygous sites on the female Z and W sequence, recoding any such sites as missing [92]. For mitochondrial genotypes we imposed only a minimum depth filter of 10x, and also masked any regions showing heteroplasmy, resulting in 14,122 remaining sites.

For individual-level assessment of genomic variation and admixture we did further filtering, imposing a minimum minor allele frequency of 0.05 in vcftools and pruning for linkage disequilibrium (LD) in plink [93]. Our pruning procedure calculated LD between each pair of single nucleotide polymorphisms (SNPs) in 50 SNP windows, removing one SNP of each pair with an r^2^ > 0.5. The window was then shifted 5 SNPs forward before repeating the procedure.

To infer demographic history and effective population sizes for *M. cardinalis*, *M. tristrami* and the outgroup *M. pulchella*, we used individuals sampled in allopatry to construct a pairwise sequential Markovian coalescent (PSMC) model [66]. We first generated a consensus sequence in fastq format for each individual using samtools mpileup and bcftools call [88], followed by limiting to autosomes and using the vcf2fq in the vcfutil.pl of bcftools. We then ran PSMC using default settings (https://github.com/lh3/psmc), and plotted output in the R v4.1.1 [94] package ggplot2 [95], using generation length of 2.37, 2.25, and 2.51 years for *M. pulchella*, *M. cardinalis*, and *M. tristrami* respectively from [96], and a per generation mutation rate of 4.6*10^-9^ from [97].

For all analyses we separated results for autosomes, neo-PAR, Z/neo-Z, W/neo-W, and the mitochondrial genome. For windowed estimates we further parsed ancestral sex chromosomes from neo-sex chromosomes. We used the program pixy [98] and an allsites VCF filtered for quality and depth to calculate nucleotide diversity (π), absolute divergence (*d*_xy_) and pairwise genetic differentiation (*F*_ST_) across 50kb windows for each population for phenotypically *cardinalis* and *tristrami* individuals. Using a quality and depth filtered VCF containing only variant sites, we used vcftools [91] to calculate Tajima’s D in 50kb windows. For all windowed analyses we limited our calculation of average estimates to 50kb windows containing a minimum of 10,000 genotyped sites, with the exception of estimates for mitochondrial sequence (which is a single window).

To quantify the degree of admixture across the genome we used the quality and depth filtered dataset to conduct ABBA-BABA tests in Dsuite [68,69]. We calculated *D* statistics for autosomes, neo-PAR, Z, neo-Z, and W/neo-W and used a block-jackknife procedure to determine if the *D* statistic is significantly different from zero. To quantify the proportion of introgression for each region we also calculated *f_4_* admixture ratio, which compares the observed excess of ABBA sites to the expected value if admixture was complete, by substituting a subset of P3 individuals for P2 individuals. The value of *f_4_* therefore provides an estimate of the proportion of admixture assuming unidirectional introgression from P3 into P2 [69,99]. Because we have allopatric populations for both parental species, we can construct and analyze two topologies to compare amount and direction of introgression between sympatric *cardinalis* and *tristrami.* The *cardinalis* P3 topology places allopatric and sympatric *tristrami* as P1 and P2 respectively and sympatric *cardinalis* as P3, therefore estimating gene flow from *cardinalis* into *tristrami.* The *tristrami* P3 topology places allopatric and sympatric *cardinalis* as P1 and P2 respectively, and sympatric *tristrami* as P3, giving an estimate of gene flow from *tristrami* into *cardinalis*. In addition to calculating *D* and *f_4_* for all autosomes, we also calculated these for each chromosome to further visualize patterns of introgression across the genome.

To understand genomic variation at an individual level and assess degree of admixture we conducted a principal components analysis (PCA) using the package SNPRelate [100] in R to investigate genomic variation of autosomes, Z/neo-Z, and W/neo-W among individuals sampled on Makira, Ugi and Three Sisters. For characterization of individuals as F1 or advanced generation backcross in the triangle plot we first identified 2,449 autosomal and 37,613 Z/neo-Z SNPs fixed between species (*F*_ST *=*_ 1) using allopatric individuals in vcftools. Using only the fixed SNPs we then calculated interspecific heterozygosity and hybrid index on the autosomes for all sympatric individuals and on the Z/neo-Z using only sympatric males in the package introgress in R [101]. Finally, we used ADMIXTURE [70] which estimates a maximum likelihood proportion of ancestry per individual and uses cross-validation error to determine the optimal number of populations or groups (K) present in sample of individuals. We ran replicate analyses using values of K from 1 to 7, and performed fivefold cross-validation to estimate error associated with each value of K.

We generated a mitochondrial haplotype network using variant sites from the mitochondrial sequence filtered for quality, minimum depth, and heteroplasmy. We used vcf2phylip [102] to convert the VCF to a phylip file for importing into the program PopART [103], where we constructed a TCS network [104] to visualize haplotype sharing and mutations separating all individuals in our dataset (including phenotypic hybrids and the outgroup *M. pulchella*).

To assess the number of private alleles for each species and location we used a custom perl script (https://github.com/ehshogren/MyzomelaPopulationGenomics) which identified monomorphic or biallelic sites that were found only in the species under consideration, present in at least five individuals of the parental species phenotype and reported which location(s) the individuals carrying that private allele were sampled from using a key file. To identify the number of variant sites with fixed differences and shared polymorphisms we used another custom perl script which considered two groups of phenotypically parental individuals (from different species and/or locations) and required a minimum of 5 individuals in each group with a genotype at the site in question.

## Acknowledgements

We thank the Solomon Islands Ministries of Environment and Education for granting research permits to work in the Solomon Islands. We also thank John and Joyce Murray, J. Tauni, J. Waihuru, K. Rupen, H. Ha’aina, J. Pepare, L. Taka, P. Teo, J. Suafuria, G. Wabeasi, and J. Waihuru for invaluable assistance in data collection.

Fieldwork was funded by the Aresty Chair in Tropical Ecology and an NSF CAREER award (IOS 1137624/0643606) to JACU, and awards from the Society of Systematic Biologists, Kushlan Graduate Research Support Fund, and Jay M. Savage Graduate Research Support Fund to JMS. Additional support provided by NSF Postdoctoral Research Fellowship to EHS (2010748), and NSF-DEB 2112474 to JACU and DCP

## Supporting Information captions

**S1 Table.**
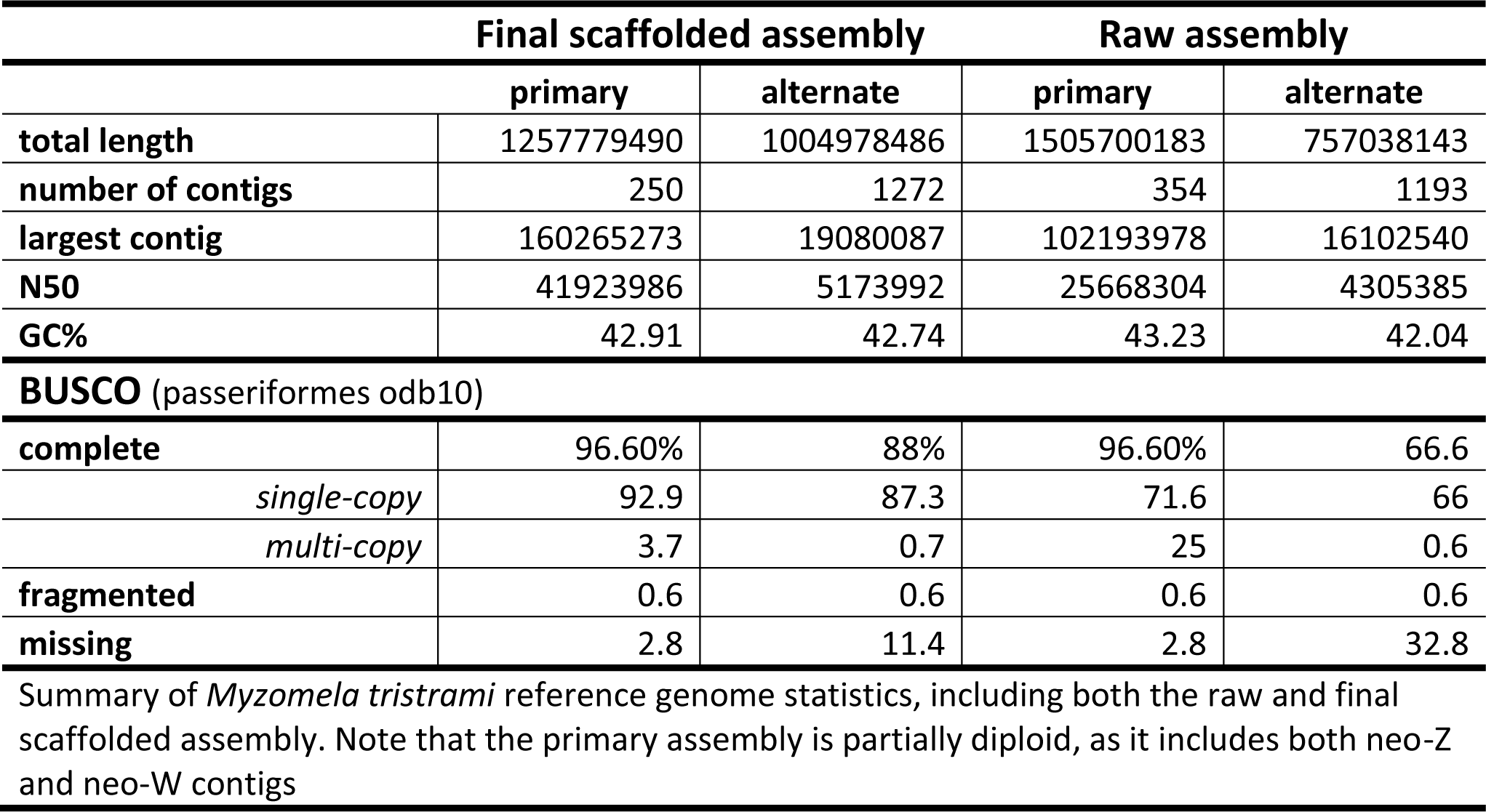
Reference genome summary. Summary of *Myzomela tristrami* reference genome statistics, including both the raw and final scaffolded assembly.

**S2 Table.** Whole genome sampling. Number of whole genome sequences by species, sampling site, and sex.

| Species | Site | Male | Female |
| --- | --- | --- | --- |
| *Myzomela cardinalis* | Ugi | 9 | 11 |
|  | Three Sisters | 10 | 10 |
|  | Sympatry (Makira) | 20 | 20 |
| *Myzomela tristrami* | Allopatry | 10 | 10 |
|  | Sympatry (Makira) | 15 | 15 |
| Phenotypic hybrid | Sympatry (Makira) | 8 | 4 |
| *Myzomela pulchella* | New Ireland |  | 1 |
| Number of whole genome sequences by species, sampling site, and sex. The *M. pulchella* sample is KU specimen catalog # 121498, KU tissue sample # 27751. |  |  |  |

**S3 Table.** SNPs per genomic region. Number of single nucleotide polymorphisms per genomic region for different filtering of datasets used in analyses.

| filtered dataset | autosome | neo-PAR | Z | neo-Z | W/neo-W | mtDNA |
| --- | --- | --- | --- | --- | --- | --- |
| Quality + depth | 27,517,646 | 601,181 | 1,431,349 | 641,775 | 90,926 | 1060 |
|  | **autosome** |  | **Z/neo-Z** |  |  |  |
| Quality + depth +<br>maf > 0.05 + LD pruned | 2,655,867 |  | 114,311 |  | 1,915 | NA |
| Number of single nucleotide polymorphisms per genomic region for different filtering of datasets used in analyses. |  |  |  |  |  |  |

**S4 Table.** Nucleotide diversity, allopatric *cardinalis* sampling sites split. Nucleotide diversity (π) averaged across 50 kb windows, standard error in parentheses for each sampled population of phenotypic parental *Myzomela cardinalis* and *M. tristrami*.

| population | autosome | neo-PAR | Z | neo-Z | W | neo-W | mtDNA |
| --- | --- | --- | --- | --- | --- | --- | --- |
| *Myzomela cardinalis* |  |  |  |  |  |  |  |
| Ugi | 0.002398<br>(7e-06) | 0.002675<br>(3.9e-05) | 0.001076<br>(2e-05) | 0.001076<br>(2.4e-05) | 3e-06<br>(1e-06) | 3e-06<br>(0) | 0.000233<br>(NA) |
| Three Sisters | 0.00198<br>(7e-06) | 0.002135<br>(4.1e-05) | 0.000588<br>(1.6e-05) | 0.000356<br>(1.4e-05) | 6e-06<br>(1e-06) | 5e-06<br>(0) | 0.000444<br>(NA) |
| Sympatry | 0.002468<br>(7e-06) | 0.002617<br>(4.1e-05) | 0.001008<br>(1.9e-05) | 0.000711<br>(2.1e-05) | 5e-06<br>(1e-06) | 4e-06<br>(0) | 0.000235<br>(NA) |
| *Myzomela tristrami* |  |  |  |  |  |  |  |
| Allopatry | 0.002759<br>(9e-06) | 0.00336<br>(4.2e-05) | 0.001738<br>(2e-05) | 0.001458<br>(1.7e-05) | 2.3e-05<br>(2e-06) | 2e-05<br>(0) | 0.001253<br>(NA) |
| Sympatry | 0.002871<br>(9e-06) | 0.003439<br>(4.1e-05) | 0.001765<br>(2.1e-05) | 0.001474<br>(1.7e-05) | 0.000587<br>(1.5e-05) | 0.000557<br>(4e-06) | 0.013763<br>(NA) |
| Nucleotide diversity ($\pi$) averaged across 50 kb windows, standard error in parentheses for each sampled population of phenotypic parental *Myzomela cardinalis* and *M. tristrami*. | | | | | | | |

**S5 Table.** Nucleotide diversity ratios. Ratio of nucleotide diversity for sex chromosome regions to large autosomes (chr1 – 10), and for the new pseudo-autosomal region (neoPAR) to a comparatively sized autosome (chr 14). Nucleotide diversity averaged across 50kb windows.

| population | Chr1 -10 $\pi$<br>(A) | Chr14 $\pi$ | neo-PAR/A | neo-PAR/<br>chr14 | Z/A | neo-Z/A | W/A | neo-W/A |
| --- | --- | --- | --- | --- | --- | --- | --- | --- |
| *Myzomela cardinalis* |  |  |  |  |  |  |  |  |
| Ugi | 0.00238 | 0.00249 | 1.124 | 1.074 | 0.452 | 0.452 | 0.001 | 0.001 |
| Three Sisters | 0.00197 | 0.00204 | 1.084 | 1.047 | 0.298 | 0.181 | 0.003 | 0.003 |
| Sympatry | 0.00245 | 0.00263 | 1.068 | 0.995 | 0.411 | 0.290 | 0.002 | 0.002 |
| *Myzomela tristrami* |  |  |  |  |  |  |  |  |
| Allopatry | 0.00269 | 0.00302 | 1.249 | 1.113 | 0.646 | 0.542 | 0.009 | 0.007 |
| Sympatry | 0.00281 | 0.00307 | 1.224 | 1.120 | 0.628 | 0.525 | 0.209 | 0.198 |

**S6 Table.** Private alleles. Number of alleles per genomic region private to each population of the respective focal species, using SNPs filtered for depth and quality only.

| population | autosome | neo-PAR | Z | neo-Z | W | neo-W | mtDNA |
| --- | --- | --- | --- | --- | --- | --- | --- |
| *Myzomela cardinalis* |  |  |  |  |  |  |  |
| Ugi | 214660 | 3809 | 12322 | 4861 | 49 | 363 | 17 |
| Three Sisters | 92008 | 1769 | 4839 | 1776 | 34 | 223 | 5 |
| Sympatry | 45480 | 701 | 2255 | 554 | 3 | 0 | 1 |
| *Myzomela tristrami* |  |  |  |  |  |  |  |
| Allopatry | 2406437 | 54121 | 143647 | 59560 | 190 | 1430 | 44 |
| Sympatry | 425142 | 11030 | 30210 | 11704 | 36 | 265 | 4 |

**S7 Table.** Fixed and shared alleles. Number of fixed and shared alleles in each region of the genome for population comparisons, using SNPs filtered for depth and quality only. Allopatry and sympatry abbreviated as Allo. and Sym. respectively

| population comparison | fix/share | autosome | neo-PAR | Z | neo-Z | W | neo-W | mtDNA |
| --- | --- | --- | --- | --- | --- | --- | --- | --- |
| *Myzomela cardinalis* |  |  |  |  |  |  |  |  |
| Ugi vs. | fixed | 1 | 0 | 1 | 4 | 0 | 0 | 0 |
| Three Sisters | shared | 5267344 | 116120 | 146136 | 57381 | 14 | 111 | 2 |
| Ugi vs. | fixed | 1 | 0 | 0 | 0 | 0 | 0 | 0 |
| Sym. *cardinalis* | shared | 6486981 | 143167 | 230689 | 121498 | 27 | 204 | 4 |
| Three Sisters vs. | fixed | 1 | 0 | 8 | 0 | 0 | 0 | 0 |
| Sym. *cardinalis* | shared | 5478982 | 118124 | 147201 | 56183 | 49 | 365 | 14 |
| *Myzomela tristrami* |  |  |  |  |  |  |  |  |
| Allo. *tristrami* vs. | fixed | 0 | 0 | 0 | 0 | 0 | 0 | 0 |
| Sym. *tristrami* | shared | 10683557 | 252252 | 477570 | 203873 | 149 | 1406 | 49 |
| Heterospecific |  |  |  |  |  |  |  |  |
| Ugi vs. | fixed | 3096 | 0 | 24421 | 9320 | 5059 | 46435 | 519 |
| Allo. *tristrami* | shared | 4875384 | 115471 | 115491 | 51977 | 0 | 3 | 0 |
| Three Sisters vs. | fixed | 30293 | 523 | 58206 | 43858 | 5062 | 46315 | 518 |
| Allo. *tristrami* | shared | 3770388 | 88987 | 57178 | 14346 | 0 | 2 | 1 |
| Ugi vs. | fixed | 5 | 0 | 22139 | 8867 | 0 | 0 | 0 |
| Sym. *tristrami* | shared | 6239777 | 139770 | 136498 | 58864 | 28 | 134 | 6 |
| Three Sisters vs. | fixed | 29 | 0 | 51495 | 39846 | 0 | 0 | 0 |
| Sym. *tristrami* | shared | 5011776 | 109917 | 70380 | 16419 | 28 | 209 | 9 |
| Allo. *tristrami* vs. | fixed | 13 | 1 | 131 | 54 | 5066 | 46360 | 516 |
| Sym. *cardinalis* | shared | 6471493 | 137238 | 158534 | 63243 | 0 | 4 | 2 |
| Sym. *cardinalis* vs. | fixed | 0 | 0 | 106 | 36 | 1 | 0 | 0 |
| Sym. *tristrami* | shared | 8036690 | 163668 | 184641 | 71334 | 50 | 340 | 13 |

**S8 Table.** Divergence between populations. Divergence (*d_xy_*) averaged across 50kb windows for each region of the genome, standard error in parentheses for pairwise comparisons of phenotypic parental populations of *Myzomela cardinalis* and *M. tristrami*. Allopatry and sympatry abbreviated as Allo. and Sym. respectively

| population comparison | autosome | neo-PAR | Z | neo-Z | W | neo-W | mtDNA |
| --- | --- | --- | --- | --- | --- | --- | --- |
| *Myzomela cardinalis* |  |  |  |  |  |  |  |
| Ugi vs.<br>Three Sisters | 0.002403<br>(7e-06) | 0.00264<br>(4e-05) | 0.00105<br>(2.1e-05) | 0.000901<br>(2.4e-05) | 5e-06<br>(1e-06) | 5e-06<br>(0) | 0.000407<br>(NA) |
| Ugi vs.<br>*Sym. cardinalis* | 0.00253<br>(7e-06) | 0.002744<br>(4e-05) | 0.001138<br>(2e-05) | 0.001014<br>(2.4e-05) | 4e-06<br>(1e-06) | 4e-06<br>(0) | 0.000246<br>(NA) |
| Three Sisters vs.<br>*Sym. cardinalis* | 0.002401<br>(7e-06) | 0.002574<br>(4.1e-05) | 0.000959<br>(2e-05) | 0.000704<br>(2.6e-05) | 6e-06<br>(1e-06) | 4e-06<br>(0) | 0.000371<br>(NA) |
| *Myzomela tristrami* |  |  |  |  |  |  |  |
| Allo. *tristrami* vs.<br>*Sym. tristrami* | 0.002833<br>(9e-06) | 0.003416<br>(4.1e-05) | 0.001782<br>(2.1e-05) | 0.001492<br>(1.7e-05) | 0.000396<br>(1e-05) | 0.000376<br>(2e-06) | 0.009029<br>(NA) |
| Heterospecific |  |  |  |  |  |  |  |
| Ugi vs.<br>*Allo. tristrami* | 0.003421<br>(9e-06) | 0.003943<br>(4.2e-05) | 0.00307<br>(2.2e-05) | 0.003088<br>(2.7e-05) | 0.001416<br>(3.6e-05) | 0.001351<br>(9e-06) | 0.040159<br>(NA) |
| Three Sisters vs.<br>*Allo. tristrami* | 0.003564<br>(9e-06) | 0.004074<br>(4.1e-05) | 0.003275<br>(2.2e-05) | 0.003403<br>(2.7e-05) | 0.001419<br>(3.7e-05) | 0.001352<br>(9e-06) | 0.040208<br>(NA) |
| *Allo. tristrami* vs.<br>*Sym. cardinalis* | 0.003403<br>(9e-06) | 0.003947<br>(4.1e-05) | 0.003132<br>(2.2e-05) | 0.003185<br>(2.9e-05) | 0.001432<br>(3.8e-05) | 0.001356<br>(9e-06) | 0.040205<br>(NA) |
| Ugi vs.<br>*Sym. tristrami* | 0.003334<br>(8e-06) | 0.003859<br>(4e-05) | 0.003054<br>(2.2e-05) | 0.003083<br>(2.7e-05) | 0.00104<br>(2.7e-05) | 0.000993<br>(6e-06) | 0.032095<br>(NA) |
| Three Sisters vs.<br>*Sym. tristrami* | 0.003453<br>(8e-06) | 0.003972<br>(3.9e-05) | 0.003256<br>(2.1e-05) | 0.003396<br>(2.7e-05) | 0.001043<br>(2.7e-05) | 0.000993<br>(6e-06) | 0.032163<br>(NA) |
| *Sym. cardinalis*<br>vs.<br>*Sym. tristrami* | 0.003313<br>(8e-06) | 0.003854<br>(3.9e-05) | 0.003117<br>(2.1e-05) | 0.003176<br>(2.8e-05) | 0.001052<br>(2.8e-05) | 0.000996<br>(6e-06) | 0.032131<br>(NA) |

Divergence ( $d_{xy}$ ) averaged across 50kb windows for each region of the genome, for pairwise comparisons of phenotypic parental populations of *Myzomela cardinalis* and *M. tristrami*. Allopatry and sympatry abbreviated as Allo. and Sym., respectively, standard errors in parentheses.

**S9 Table.**
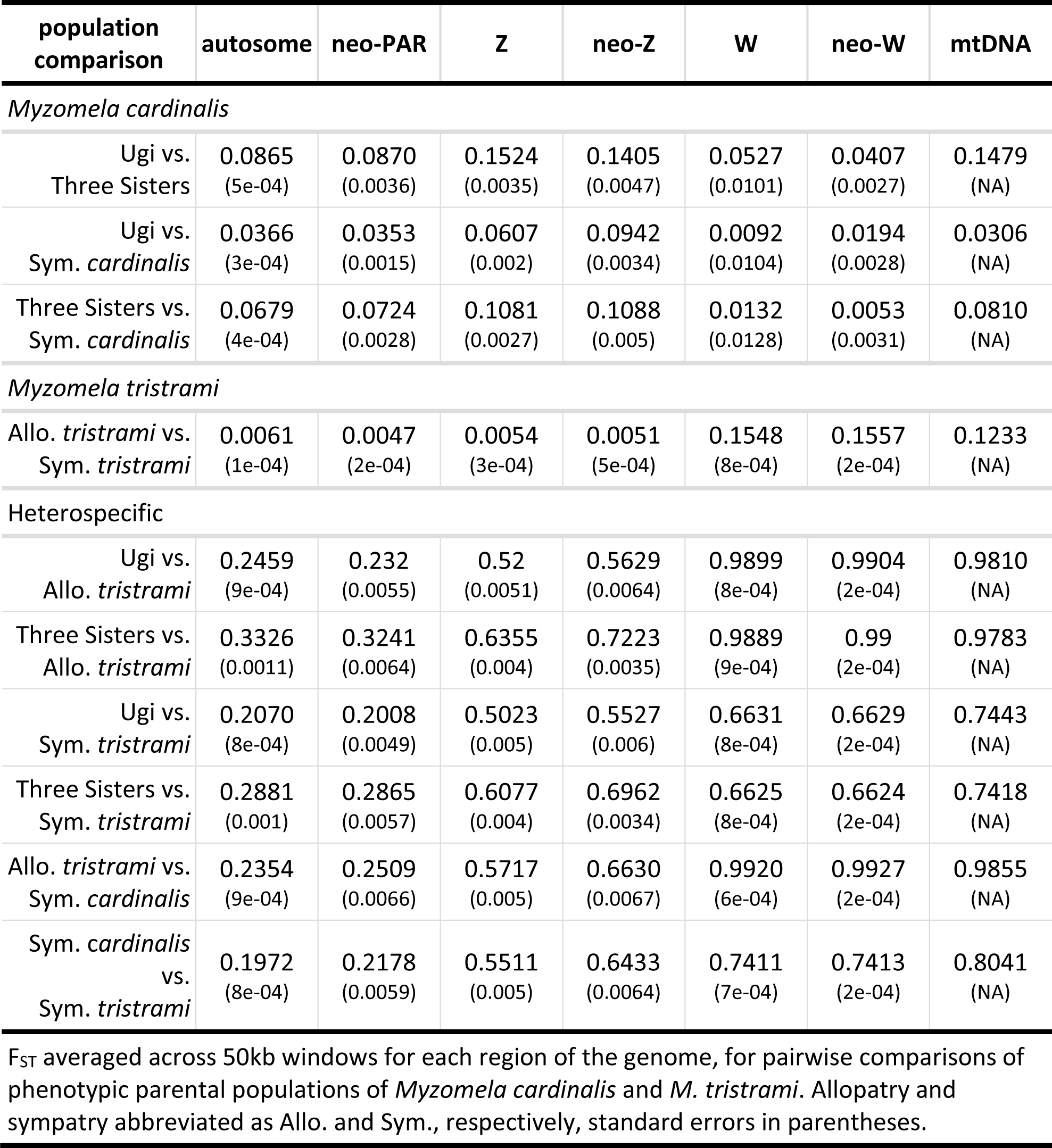
Differentiation between populations, allopatric *cardinalis* sampling sites split. Differentiation (F_ST_), averaged across 50kb windows for each region of the genome, standard error in parentheses for pairwise comparisons of phenotypic parental populations of *Myzomela cardinalis* and *M. tristrami*. Allopatry and sympatry abbreviated as Allo. and Sym. respectively.

| population comparison | autosome | neo-PAR | Z | neo-Z | W | neo-W | mtDNA |
| --- | --- | --- | --- | --- | --- | --- | --- |
| *Myzomela cardinalis* |  |  |  |  |  |  |  |
| Ugi vs.<br>Three Sisters | 0.0865<br>(5e-04) | 0.0870<br>(0.0036) | 0.1524<br>(0.0035) | 0.1405<br>(0.0047) | 0.0527<br>(0.0101) | 0.0407<br>(0.0027) | 0.1479<br>(NA) |
| Ugi vs.<br>Sym. *cardinalis* | 0.0366<br>(3e-04) | 0.0353<br>(0.0015) | 0.0607<br>(0.002) | 0.0942<br>(0.0034) | 0.0092<br>(0.0104) | 0.0194<br>(0.0028) | 0.0306<br>(NA) |
| Three Sisters vs.<br>Sym. *cardinalis* | 0.0679<br>(4e-04) | 0.0724<br>(0.0028) | 0.1081<br>(0.0027) | 0.1088<br>(0.005) | 0.0132<br>(0.0128) | 0.0053<br>(0.0031) | 0.0810<br>(NA) |
| *Myzomela tristrami* |  |  |  |  |  |  |  |
| Allo. *tristrami* vs.<br>Sym. *tristrami* | 0.0061<br>(1e-04) | 0.0047<br>(2e-04) | 0.0054<br>(3e-04) | 0.0051<br>(5e-04) | 0.1548<br>(8e-04) | 0.1557<br>(2e-04) | 0.1233<br>(NA) |
| Heterospecific |  |  |  |  |  |  |  |
| Ugi vs.<br>Allo. *tristrami* | 0.2459<br>(9e-04) | 0.232<br>(0.0055) | 0.52<br>(0.0051) | 0.5629<br>(0.0064) | 0.9899<br>(8e-04) | 0.9904<br>(2e-04) | 0.9810<br>(NA) |
| Three Sisters vs.<br>Allo. *tristrami* | 0.3326<br>(0.0011) | 0.3241<br>(0.0064) | 0.6355<br>(0.004) | 0.7223<br>(0.0035) | 0.9889<br>(9e-04) | 0.99<br>(2e-04) | 0.9783<br>(NA) |
| Ugi vs.<br>Sym. *tristrami* | 0.2070<br>(8e-04) | 0.2008<br>(0.0049) | 0.5023<br>(0.005) | 0.5527<br>(0.006) | 0.6631<br>(8e-04) | 0.6629<br>(2e-04) | 0.7443<br>(NA) |
| Three Sisters vs.<br>Sym. *tristrami* | 0.2881<br>(0.001) | 0.2865<br>(0.0057) | 0.6077<br>(0.004) | 0.6962<br>(0.0034) | 0.6625<br>(8e-04) | 0.6624<br>(2e-04) | 0.7418<br>(NA) |
| Allo. *tristrami* vs.<br>Sym. *cardinalis* | 0.2354<br>(9e-04) | 0.2509<br>(0.0066) | 0.5717<br>(0.005) | 0.6630<br>(0.0067) | 0.9920<br>(6e-04) | 0.9927<br>(2e-04) | 0.9855<br>(NA) |
| Sym. *cardinalis*<br>vs.<br>Sym. *tristrami* | 0.1972<br>(8e-04) | 0.2178<br>(0.0059) | 0.5511<br>(0.005) | 0.6433<br>(0.0064) | 0.7411<br>(7e-04) | 0.7413<br>(2e-04) | 0.8041<br>(NA) |

$F_{ST}$ averaged across 50kb windows for each region of the genome, for pairwise comparisons of phenotypic parental populations of *Myzomela cardinalis* and *M. tristrami*. Allopatry and sympatry abbreviated as Allo. and Sym., respectively, standard errors in parentheses.

**S10 Figure.**
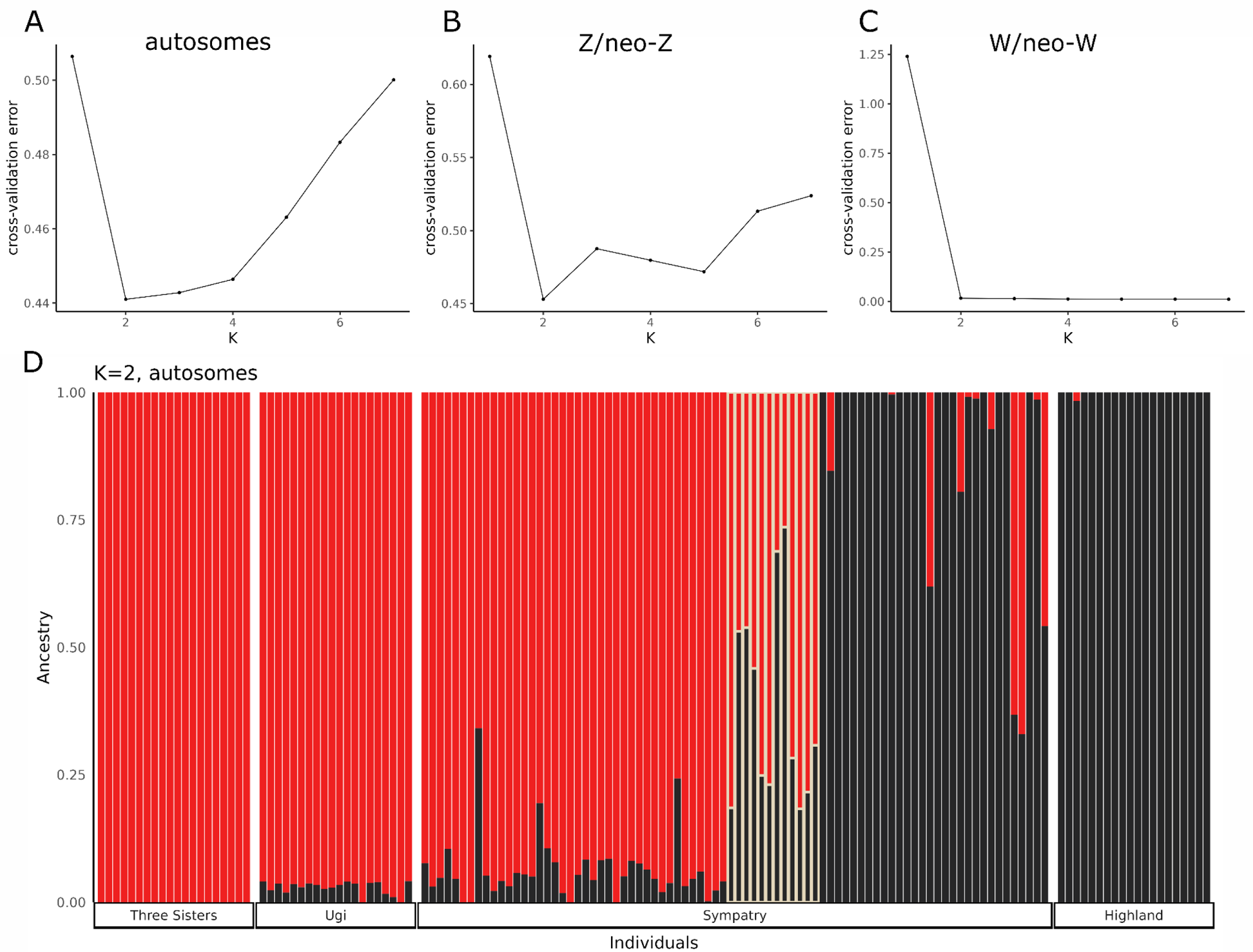
ADMIXTURE cross-validation error and plot of autosomes at K=2. Cross-validation error for ADMIXTURE of K = 1-7 for autosomes (A), Z/neo-Z (B), and W/neo-W (C). ADMIXTURE plot for autosomal sequence with K = 2 (D). Phenotypic hybrids are outlined in yellow, with phenotypic *cardinalis* to the left and phenotypic *tristrami* to the right of phenotypic hybrids.

**S11 Figure.**
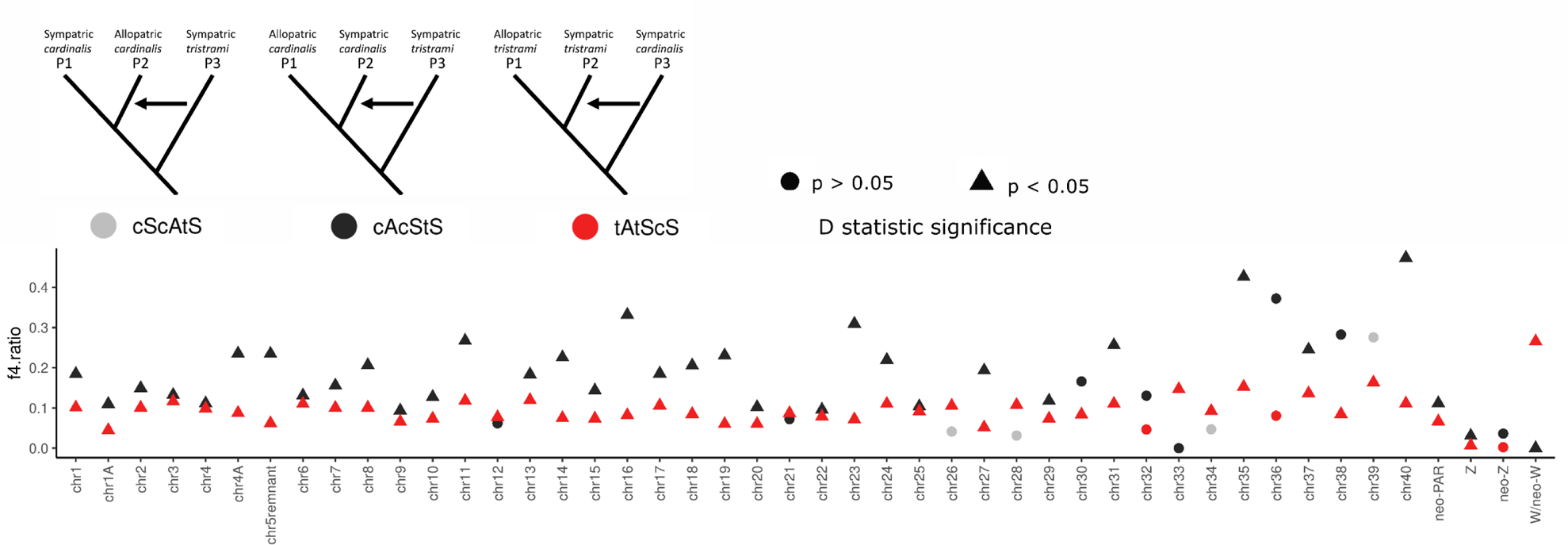
*f_4_* admixture ratio, all topologies. Admixture ratio (*f_4_* statistic) for each autosome and sex chromosome regions. Color of the point indicates for which topology the statistic was calculated, and shape of the point indicates whether the *D* statistic for that chromosome was significantly different from zero, using the block-jackknife procedure.

